# Bacterial family-specific enrichment and functions of secretion systems in the rhizosphere

**DOI:** 10.1101/2024.05.07.592589

**Authors:** A. Fourie, J.L. Lopez, J.J. Sánchez-Gil, S.W.M. Poppeliers, R. de Jonge, B.E. Dutilh

**Affiliations:** Theoretical Biology and Bioinformatics, Department of Biology, Science for Life, Utrecht University, 3584CH, Utrecht, The Netherlands; Instituto de Biotecnología y Biología Molecular, CONICET, CCT-La Plata, Departamento de Ciencias Biológicas, Facultad de Ciencias Exactas, Universidad Nacional de La Plata, La Plata, Argentina; Plant-Microbe Interactions, Department of Biology, Science for Life, Utrecht University, 3584CH, Utrecht, The Netherlands; AI Technology For Life, Department of Information and Computing Sciences, Science for Life, Utrecht University, 3584CC, Utrecht, The Netherlands; Institute of Biodiversity, Faculty of Biological Sciences, Cluster of Excellence Balance of the Microverse, Friedrich-Schiller-University Jena, 07743, Germany

## Abstract

The plant rhizosphere is a highly selective environment where bacteria have developed traits to establish themselves or outcompete other microbes. These traits include bacterial secretion systems (SS’s) that range from Type I (T1SS) to Type IX (T9SS) and can play diverse roles. The best known functions are to secrete various proteins or other compounds into the extracellular space or into neighbouring cells, including toxins to attack other microbes or effectors to suppress plant host immune responses. Here, we aimed to determine which bacterial SS’s were associated with the plant rhizosphere. We utilised paired metagenomic datasets of rhizosphere and bulk soil samples from five different plant species grown in a wide variety of soil types, amounting to ten different studies. The T3SS and T6SS were generally enriched in the rhizosphere, as observed in studies of individual plant-associated genera. We also identified additional SS’s that have received less attention thus far, such as the T2SS, T5SS and *Bacteroidetes*-specific T6SSiii and T9SS. The predicted secreted proteins of some of these systems (T3SS, T5SS and T6SS) could be linked to functions such as toxin secretion, adhesion to the host and facilitation of plant-host interactions (such as root penetration). The most prominent bacterial taxa with rhizosphere- or soil-enriched SS’s included *Xanthomonadaceae*, *Oxalobacteraceae*, *Comamonadaceae*, *Caulobacteraceae*, and *Chitinophagaceae,* broadening the scope of known plant-associated taxa that use these systems. We anticipate that the SS’s and taxa identified in this study may be utilised for the optimisation of bioinoculants to improve plant productivity.

## Introduction

Many microbes have symbiotic relationships with plants and are able to colonise the above and below-ground parts of the plant. These plant-associated microbes can act as pathogens, commensals or beneficials. A strong focus for the development of sustainable agriculture has been on utilising plant growth-promoting rhizobacteria (Rosier *et al*., 2018, Lucke *et al*., 2020) and biocontrol agents that can contribute directly to plant growth by producing phytohormones (Finkel *et al*., 2020), facilitating nutrient uptake such as iron, or by producing antibiotics to fend off pathogens (Bakker *et al*., 2020, Pascale *et al*., 2020). It is thus essential to understand which microbes are beneficial, how they function and how they can be optimised for agricultural applications (Poppeliers *et al*., 2023).

The root environment is a highly selective environment, influenced by plant root exudates (Pascale *et al*., 2020). This results in a reduced taxonomic diversity closer to the root surface, referred to as the rhizosphere effect (Bakker *et al*., 2013, Schneijderberg *et al*., 2020). Near to the plant root, fast-growing microbes are selected (López *et al*., 2023) that can compete for nutrients and for attachment to the host surface. Other traits that allow bacteria to colonise and persist in the rhizosphere include flagellar motility (Li *et al*., 2021, López *et al*., 2023, Sánchez-Gil *et al*., 2023), attachment and biofilm formation (Seneviratne *et al*., 2011), processing of plant metabolites (Barret *et al*., 2011, Sánchez-Gil *et al*., 2023) and iron chelation (Lemanceau *et al*., 2009).

Bacterial secretion systems (SS’s) represent another trait that likely contributes to competition and colonisation in the rhizosphere. These systems include membrane-spanning protein channels that facilitate secretion or translocation of proteins or other molecules that contribute to extracellular nutrient processing, function as effector proteins for host interactions, or as toxins to attack neighbouring microbes (Bernal *et al*., 2018, Lucke *et al*., 2020). Known SS’s are widely distributed among gram-negative bacteria (Abby *et al*., 2016, Green *et al*., 2016), they are prominent among *Proteobacteria* and some other gram-negative phyla and are classified into nine main types, ranging from Type 1 (T1SS) to Type 9 (T9SS). One sub-type of the T6SS, T6SSiii, and the T9SS are restricted to *Bacteroidetes* (James *et al*., 2022), and a specific T7SS, known as the ESAT-6 SS, is only observed in some genera of the gram-positive *Firmicutes* and *Actinobacteria* phyla (Abdallah *et al*., 2007, Ates *et al*., 2016). SS’s can be associated to different membranes. In gram negative bacteria, some SS channels are located only in the outer membrane (T2SS, T5SS) while others cross both membranes and some can even penetrate neighbouring bacterial or eukaryotic cells (T3SS, T4SS and T6SS).

Distinct functions have been associated with different SS types, based on their secreted proteins. For example, T1SS’s contribute to antagonism or host attachment by secretion of toxins or adhesins (Thomas *et al*., 2014, Hui *et al*., 2021), T5aSS’s and T5cSS’s are mostly linked to membrane attachment (Fan *et al*., 2016) and T5bSS’s are involved in contact-dependent growth inhibition (CDI) for interbacterial competition (Willett *et al*., 2015). T3SS’s can suppress host immune responses through translocation of effector proteins (Zboralski *et al*., 2022) or by contributing to biofilm formation (Castiblanco & Sundin, 2016), and T6SS’s secrete toxic effectors intracellularly to attack competing microbes (Gallegos-Monterrosa & Coulthurst, 2021).

Functional studies of specific plant-associated bacterial genera support the importance of SS’s (including T3SS, T6SS and T7SS) in rhizosphere colonization and survival. In *Pseudomonas,* the T6SS was shown to contribute to interbacterial competition and persistence in the rhizosphere (Bernal *et al*., 2017, Durán *et al*., 2021, Boak *et al*., 2022), in *Rhizobium* it played a role in biocontrol (Vogel *et al*., 2021) and in *Xanthomonadales* it contributed to antagonism against prokaryotes and eukaryotes (Bayer-Santos *et al*., 2019). The T3SS has been shown to facilitate root colonisation of legume-associated *Burkholderia* species (Wallner *et al*., 2021) and for plant growth-promoting *Pseudomonas* strains it can contribute to biocontrol properties or suppression of plant immune responses (Zboralski *et al*., 2022). Plant-beneficial *Bacilli* utilise T7SS’s to secrete a protein that embeds into the plant plasma membrane and causes iron leakage, facilitating root colonisation (Liu *et al*., 2023).

Various community-based metagenomic analyses have reported an enrichment in the rhizosphere (vs bulk soil) of some bacterial SS proteins. For example, in cucumber and wheat various genes from the T2SS, T3SS and T6SS were enriched (Ofek-Lalzar *et al*., 2014), and in barley genes from the T3SS and T6SS were enriched (Bulgarelli *et al*., 2015). In genome-based comparisons of plant-associated and soil-isolated microbes, T3SS’s and T6SS’s were also enriched in plant-associated microbes (Levy *et al*., 2018) and specifically in copiotrophs, fast-growing microbes in nutrient-rich environments such as the rhizosphere (López *et al*., 2023). However, these studies evaluated the different protein components of a SS independently and not as a unit. Except in the case of the T5SS, SS proteins rarely function alone but rather as complexes whose genes occur as operons in the genome. Another challenge for SS analysis is that some proteins can also form part of more than one SS. For example, secretin forms part of the T2SS and the T3SS (Denise *et al*., 2020) and the gene content of the T6SS and the extracellular contractile injection system (eCIS) are highly similar (Geller *et al*., 2021). Such functional redundancy can confound abundance estimates of SS’s and should be taken into account when interpreting results.

To our knowledge, there has not been a comprehensive investigation of all bacterial SS’s present in natural root communities of diverse plant species. In addition, the approach of considering all protein components of a SS as a unit to confirm its presence, while excluding redundant proteins, has not been used before, but could impact the detection of prominent SS’s. The aim of this study was to determine which SS’s were enriched in the rhizosphere environment in comparison to unplanted soil, as an indication of their involvement in rhizosphere colonisation and survival. Publicly available metagenome sequencing studies, that evaluated paired rhizosphere and soil samples, were investigated for the presence and abundance of all well-defined bacterial SS’s (T1SS-T6SS and T9SS). The studied pairs included diverse plants, i.e. *Arabidopsis thaliana* (two studies), cucumber, wheat, chickpea and various citrus species, allowing us to investigate consistencies between different plant species. In addition to quantifying the SS’s, we also assessed their association to bacterial families. Finally, we predicted the secreted proteins of the SS’s of interest, including the functional annotation of effector proteins, as an indication of the functional role of the SS’s and their contribution to their host bacteria.

## Results

### Metagenomic data processing

In this study we utilised existing paired metagenomic datasets of rhizosphere and soil samples, from various plant species (Table S1). This included *Arabidopsis thaliana* (Sanchez-Gil *et al*., Unpublished, Stringlis *et al*., 2018), chickpea and wheat (Zhou *et al*., 2020), citrus species from Brazil, Italy, China and Spain (Xu *et al*., 2018), and cucumber and wheat (Ofek-Lalzar *et al*., 2014). The number of raw sequencing reads per dataset ranged from 26M to 115M reads per sample (Table S1), with the datasets from *Arabidopsis* (Stringlis *et al*., 2018), wheat and cucumber (Ofek-Lalzar *et al*., 2014) being in the lower range and citrus having the largest datasets (Xu *et al*., 2018). For ease of reference, these datasets were named as follows: Arabidopsis_Stringlis, Arabidopsis_Sanchez, Chickpea_Zhou, Wheat_Zhou, Cucumber_Ofek, Wheat_Ofek, Citrus_Brazil, Citrus_China, Citrus_Italy, Citrus_Spain. Sequences were quality filtered, host reads removed and then assembled into contigs to allow for gene and protein predictions.

The rhizosphere sampling method had a big impact on the number of reads retained after host read removal. In the chickpea and wheat datasets (Zhou *et al*., 2020) and citrus datasets (Xu *et al*., 2018) about 97% of the reads were retained as microbial reads whereas for the other studies only 50-60% of the reads were of microbial origin and the remainder were derived from the plant host (see Table S1). In the former studies the soil adhering to the roots was collected and only the soil was used for DNA isolation and sequencing, whereas the other studies included the plant root and the adhering soil. This means that likely more of the tightly-adhering microbes and endophytes were included in the latter studies.

The total assembly length varied greatly between datasets, ranging from 72 Mbp in datasets with fewer reads up to 1 Gbp in datasets with the most reads (Table S1). The N50 values of the assemblies also showed an upward trend where datasets with more reads had larger assemblies. However, the *Arabidopsis* datasets had particularly high N50 values, relative to the amount of input reads. This could be an effect of the sampling method and consequently a lower bacterial diversity present in these datasets.

The quality of the contigs’ assemblies was assessed by the percentage of reads that mapped back to contigs, thus indicating what proportion of the reads were assembled. For most of the datasets an average of 10-20% of the reads mapped to the assembled contigs (Table S1). This low percentage has also been observed in other soil microbiome assemblies and is a result of the high diversity of the soil microbiome. This suggests current algorithms and depth of coverage have a limited resolution to construct inclusive metagenomic assemblies in soil (Howe *et al*., 2014, Xu *et al*., 2021).

### Rhizosphere sampling methods impact observation of a rhizosphere effect in metagenomes

Previous studies have shown a lower microbial diversity near the root (rhizosphere effect) suggesting stronger selection for specific microbes and functions (Bakker *et al*., 2013, Schneijderberg *et al*., 2020). The metagenomic datasets used in this study used different sampling methods, as mentioned earlier. Due to these differences, we wanted to determine if a clear rhizosphere effect could still be observed in all datasets. To this end, we first performed taxonomic read classification of the metagenomic datasets using Kaiju (Menzel *et al*., 2016). The taxonomic identity could be determined for 82% (± 3.6 SD) of the metagenomic reads of each dataset. Of these, 45% (± 7.1 SD) were classified at the genus rank and 51% (± 8.8 SD) at the family rank (Fig. S1). The taxonomic composition and core genera observed in the datasets are summarised in Fig. S2. Using the taxonomic quantification of the reads, we next computed the per-sample alpha diversity at the genus rank, applying the Shannon index that quantifies the number of different organisms and their evenness. This enabled us to compare soil versus rhizosphere alpha diversity per dataset. The alpha diversity was significantly lower in the rhizosphere than in the soil for both *Arabidopsis* datasets, the Citrus_Brazil and the Wheat_Ofek datasets, but significantly higher for the Chickpea_Zhou and the Wheat_Zhou datasets (Fig. 1A; t-test, *p value* < 0.05, BH-correction). The other datasets did not show a strong rhizosphere effect either way. The rhizosphere effect was thus predominantly observed in datasets where the root and tightly adhering microbes were included.

**Figure 1.**
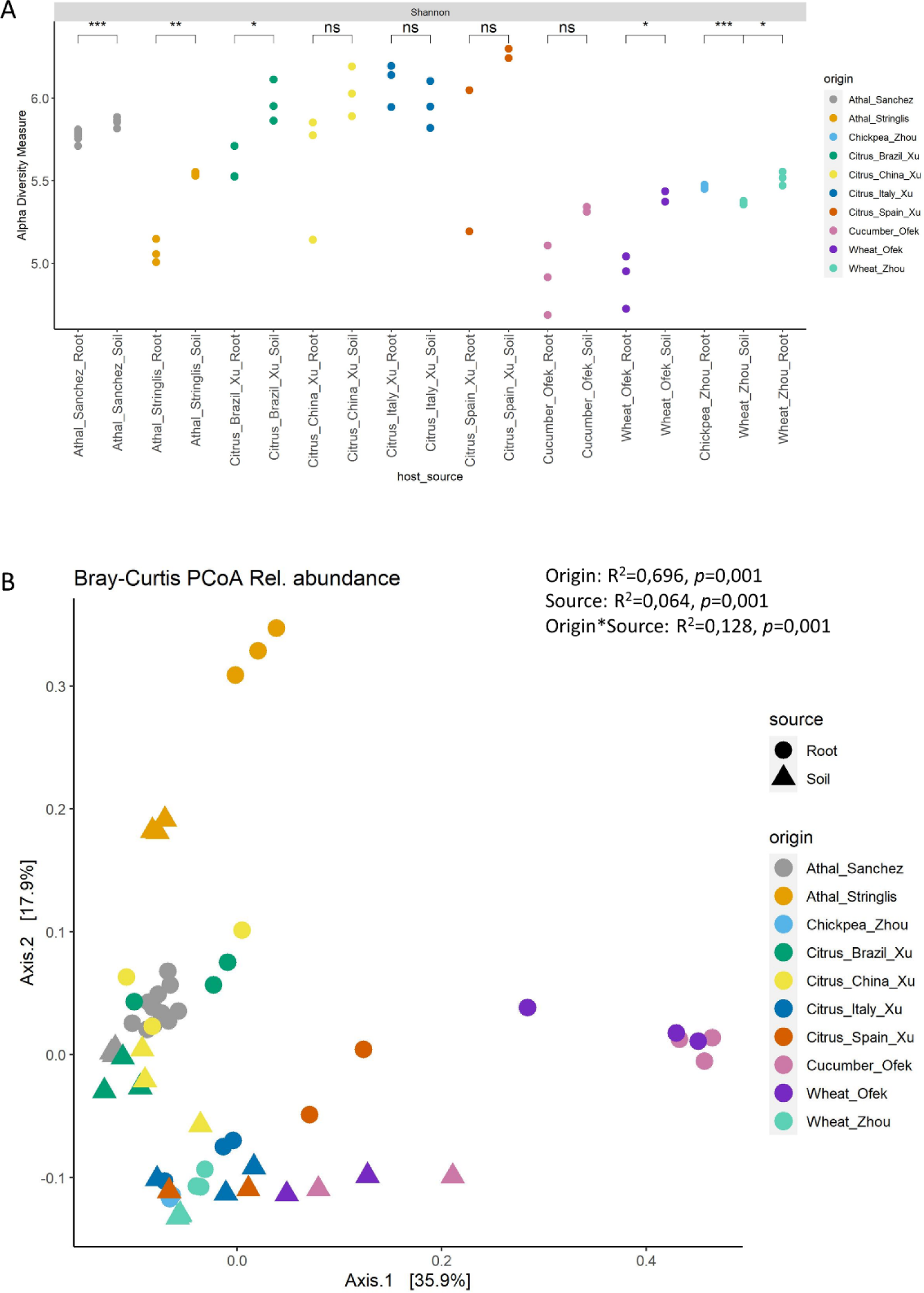
Alpha- and beta diversity of all metagenomic datasets, based on genus rank classification of metagenomic reads (Kaiju). A) The Shannon-diversity index was compared between the rhizosphere and soil samples of each plant species to determine the rhizosphere effect (lower diversity in the rhizosphere) in each study. Studies with significant differences (t-test, *p value* < 0.05, BH-correction) are indicated with an asterisk. B) Pairwise sample-to-sample Bray-Curtis dissimilarities were used to perform principal coordinate analysis (based on genera > 1·10^-5^ relative abundance) to illustrate differences in community composition between plant species and different biomes (rhizosphere vs soil).

Visualisation of the beta diversity of the different datasets in a multivariate analysis PCoA plot showed distinct clustering of root and soil samples of most datasets, except Citrus_China and Citrus_Italy (Fig. 1B; see Fig. S3 for individual plots). In the combined analysis of all datasets (Fig. 1B), the metagenomic dataset source had the biggest effect on the variation observed, but the interaction between metagenomic dataset and biome (root or soil) was also significant. This suggests some distinct differences between the root and soil communities, per dataset. However, in individual analyses of most datasets the differences were not statistically significant (permanova, *p value* >0.05; Table S2), except for the Arabidopsis_Sanchez dataset. This corresponds to the low distinction between biomes and the low rhizosphere effect observed in the alpha diversities.

### Four secretion systems showed rhizosphere enrichment in the metagenomic data

Twelve subtypes of well-defined SS’s were investigated, including the T1SS - T6SS and T9SS and their respective subtypes (T4SSI, T4SST, T5aSS, T5bSS, T5cSS, T6SSi, T6SSii and T6SSiii). To identify all possible SS structural proteins in the metagenomic datasets and not only those that occurred in assembled contigs, since only 10-20% of the reads were assembled, we used a read mapping approach (see M&M for details). To quantify all the possible SS structural proteins, we first created a comprehensive SS protein database. The first part of this database was created using all 254,090 bacterial genomes in the Genome Taxonomy Database (GTDB) (Parks *et al*., 2021), in order to represent as many taxa as possible. SS’s were detected in 129,743 genomes, representing 116 phyla divided into 1,441 families and 4,525 genera (Table S3). Due to the ambiguity of some SS proteins involved in different SS’s (see Introduction), a subset of representative proteins was selected per SS. A non-redundant GTDB protein database of 263,396 proteins was created. For each metagenomic dataset a study-specific reference protein database was constructed, consisting of the GTDB SS proteins and those identified in the assembled metagenomic contigs.

The DIAMOND blastx algorithm (Buchfink *et al*., 2015) was used to map metagenomic reads to the SS protein database. Identified proteins were filtered based on percentage sequence identity and percentage coverage of the read mappings. Mapped read counts were converted and normalised similar to transcripts per kilobase million (TPM), but to avoid confusion with RNA transcripts was termed as metagenomic reads per kilobase million (MRPM) in this manuscript.

Summaries of the read mapping results confirmed the presence of the T1SS-T3SS, T4SS_T_, T5a-cSS, T6SSi, T6SSiii and T9SS in the metagenomic datasets. Some types of SS’s were more abundant in the metagenomes than others. The T1SS was the most abundant, followed by T5aSS and T5bSS (Fig. 2). This corresponds to what was observed in the GTDB bacterial genomes (Table S3) where the T1SS, T5aSS and T5bSS were detected in the largest number of bacterial families (765, 542 and 499, respectively), illustrating that some SS’s are more widely distributed among bacterial taxa than others.

**Figure 2.**
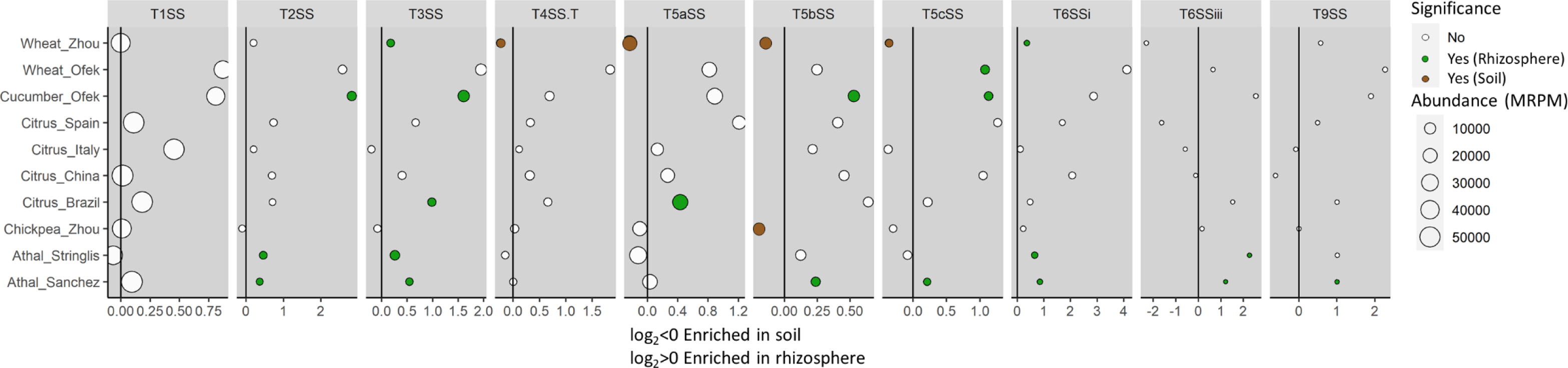
Secretion systems enriched in the rhizosphere or soil in each plant species. Green and brown circles indicate SS’s that were significantly more abundant in one of the biomes (t-test, *p value* <0.05, BH-correction). Some SS’s were more abundant in a specific biome but not significantly so, mostly due to the variation between samples in a dataset. The abundance in the rhizosphere relative to the soil is indicated on the x-axis as log-fold values (positive enriched in rhizosphere, negative enriched in soil) and circle sizes represent the MRPM abundance of the SS in a plant dataset.

The abundance (MRPM value) of the SS’s in the rhizosphere and soil biomes of the different studies were compared to evaluate if there is a higher abundance of specific SS’s in one of the two biomes. From a total of 100 comparisons (10 SS’s in 10 metagenomic datasets), 20 SS’s were significantly enriched in rhizosphere samples and 5 were enriched in soil samples. The remaining 75 comparisons were not significant. All SS’s except the T1SS showed enrichment in one or multiple plant datasets (Fig. 2). General trends could be observed across the different datasets; four SS types were more abundant in the rhizosphere in multiple plant host datasets (Fig. 2). This included the T2SS (both *Arabidopsis* sets and Cucumber_Ofek), T3SS (both *Arabidopsis* sets, Cucumber_Ofek and Wheat_Zhou), T5cSS (Arabidopsis_Sanchez, Cucumber_Ofek and Wheat_Ofek) and T6SSi (both *Arabidopsis* sets and Wheat_Zhou). The soil-enriched SS’s (T4SS_T_, T5aSS, T5bSS, T5cSS) were all found in the Zhou datasets, where we also observed an unexpected reverse rhizosphere effect (Fig. 1a). In the latter dataset, the soil was collected from a farm site where different crops have been rotated and soil management included no tillage. This could impact the existing microbial community and perhaps explain the contrasting results observed.

### Core bacterial families can be linked to the identified SS’s

Based on the Kaiju classification of the metagenomic reads, a total of 559 bacterial families could be identified in the investigated metagenomic datasets. To identify which families in the rhizosphere utilise SS’s, and to estimate their abundance, the identified SS proteins were also taxonomically classified up to family rank (see M&M). The number of families in which the different SS’s were detected ranged from 267 families (48%) with a T1SS, to 104 – 170 (19% - 30%) with a T3SS, T5SS or T6SSi, to low numbers of 4 (0.7%) and 22 (4%) families with a T6SSiii and T9SS, respectively.

The families in which SS’s were detected in the majority of the plant datasets (≥80%) are of specific interest. These can be considered core families whose members utilise the SS’s in different rhizosphere environments. A total of 52 core bacterial families were identified among the metagenomic datasets (Fig. 3). The T1SS was found in the largest number of core bacterial families (37 families), while *Chitinophagaceae* was the only core family in which a T6SSiii and T9SS were consistently detected in all plant studies. Other SS’s of interest included the T2SS which was consistent in 10 bacterial families across the plant studies, the T3SS which was found in 18 families and the T6SSi which was identified in 15 families. Since most community-based metagenomic studies primarily focused on functional enrichment and not taxonomic composition of the SS’s, this study adds new knowledge on the types of families that may encode for SS’s in the rhizosphere. For example, the T3SS’s identified in the *Solirubrobacteraceae*, *Bryobacteraceae*, *Oxalobacteraceae* and *Caulobacteraceae* families and the T5bSS’s which have, to our knowledge, primarily been identified in plant-associated *Pseudomonas* (Berendsen *et al*., 2015), *Rhizobium* (Liang *et al*., 2018) and *Paraburkholderia* (Dias *et al*., 2019) species.

**Figure 3.**
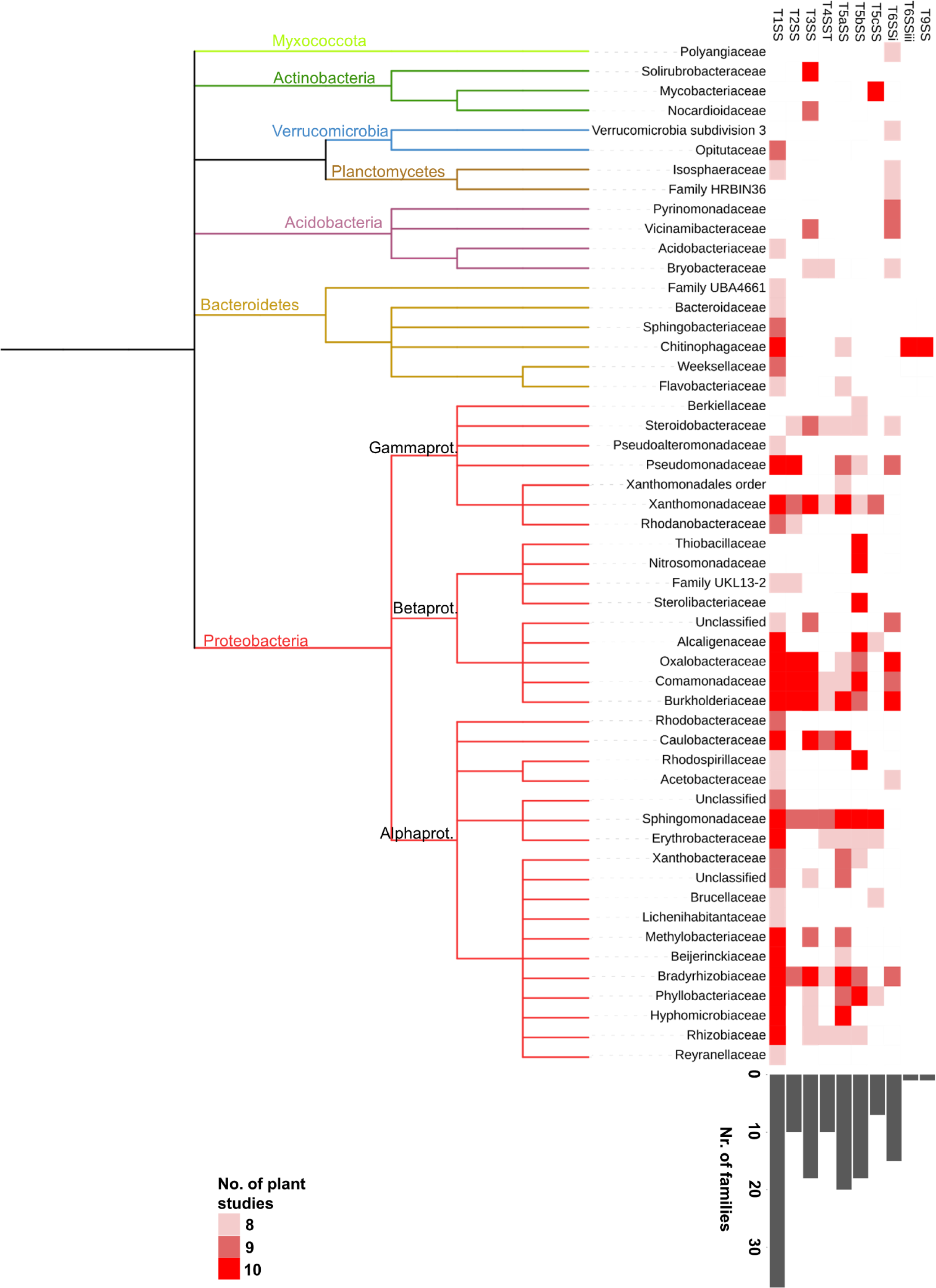
Fifty-two bacterial families encoding SS’s in the rhizosphere (detected in ≥80% of the plant studies/datasets). The coloured gradient boxes relate to the number of plant studies in which a family with the corresponding SS was detected (8-10 studies). Some families were detected in all studies considered (dark red). The bar plot at the bottom summarises the total number of families in which a SS was observed.

By comparing the total abundance of the different SSs between rhizosphere and soil, many of the SS’s were not clearly enriched in either biome (Fig. 2). However, investigating SS abundance per family revealed specific families with enriched SS’s in the rhizosphere or soil. A total of 41 families with SS’s showed enrichment in multiple plant species (Fig. 4A). If the SS’s of the same family are enriched in different plant rhizosphere environments, this suggests it is a common trait in this family and likely plays an important role for some of its species. These families were thus of specific interest.

**Figure 4.**
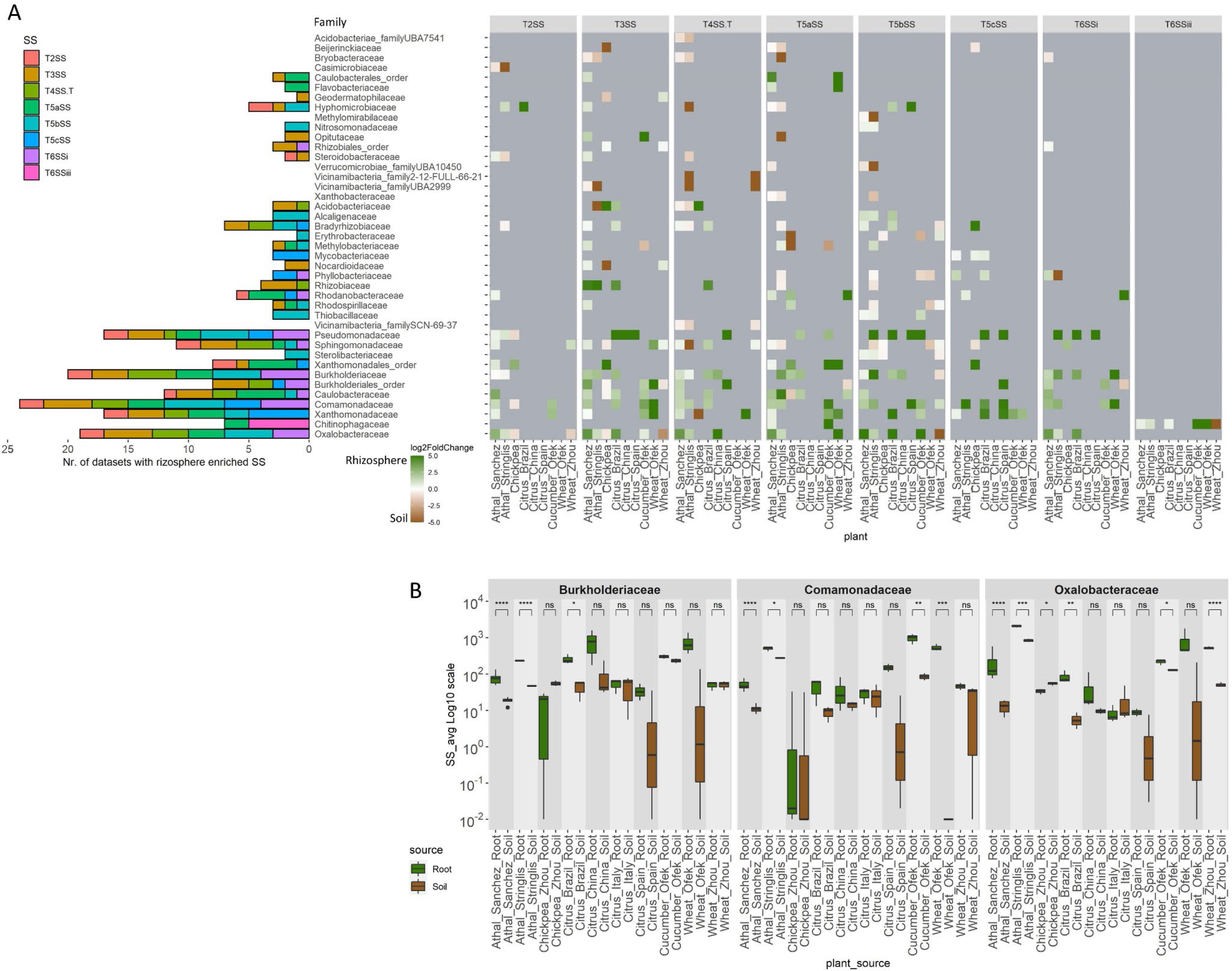
Enrichment of SS’s of specific families in all plant studies/datasets investigated. A) The log2fold difference ranges from green (enriched in the rhizosphere) to brown (enriched in the soil). Families that showed a significant enrichment in rhizosphere or soil for a specific SS in more than one dataset is displayed, starting with the families present in most datasets from the bottom of the figure. The counts of SS’s per family is summarised on the left in a barplot. B) Three prominent families, *Oxalobacteriaceae, Burkholderiales* and *Comamonadaceae,* showed enrichment of the T3SS in multiple datasets. The log10(MRPM abundance) of the T3SS is shown for the rhizosphere and soil biomes across the different studies and plants.

One prominent family was the *Oxalobacteraceae* that was enriched in the rhizosphere in multiple plant datasets for the T3SS, T4SST, T5aSS and the T6SS (Fig. 4B). Other families of interest were *Burkholderiaceae* (T3SS, T5SS, T6SS) (Fig. 4B), *Comamonadaceae* (T3SS, T4SST, T5SS and T6SSi) (Fig. 4B), *Xanthomonadaceae* (T3SS, T5cSS), *Caulobacteraceae* (T3SS, T5SS and T6SS), *Pseudomonadaceae* (T3SS, T5cSS and T6SS) and some unclassified families in the *Burkholderiales* order. In five of the studies the *Chitinophagaceae* also showed an enrichment of the T6SSiii in the rhizosphere (Fig. 4A).

### Functional prediction of SS’s in prominent bacterial families - *Arabidopsis thaliana* as model

The aforementioned metagenomic data analyses suggested that SS’s were especially important for some of the microbes in the rhizosphere. Next, we focused on predicting the functional roles of these SS’s and the potential presence of multiple copies of a SS within a genome. This information cannot be reliably predicted from the metagenomic data, so we constructed a genomic dataset of microbes isolated from the plant rhizosphere. A large collection of 241 genomes is publicly available for *A. thaliana*-associated microbes (referred to as At_microbiome genomes, Table S4, see M&M) and further investigations were thus narrowed to this host. This also allowed identification of the most prominent SS-encoding genera within the families.

### The proteins secreted by the T3SS, T5SS and T6SSi facilitate diverse plant and microbe interactions

The prediction of secreted products has most reliably been established for the T3SS, T4SS, T5SS and T6SS (Gerlach & Hensel, 2007, An *et al*., 2016, Fan *et al*., 2016, Zeng & Zou, 2017) and predictions were thus restricted to these SS’s. The T4SS was excluded, as we focused on those that showed enrichment. The T3SS, T4SS and T6SS directly inject effector proteins into a host cell to interact with host cellular proteins (Galán, 2009), whereas T5SS’s also secrete other proteins extracellularly, such as proteases or adhesins. Secreted products were either predicted based on surrounding structural SS genes in the genome, or by blastp comparisons of the proteome to known effectors (for details see M&M).

Of the 241 genomes analysed, 62 encoded either one or two copies of a T3SS (Table S5). Based on blastp comparisons of the proteomes to known effectors, a total of 339 putative effectors were identified in these genomes, with a range of 1-13 effectors per genome (average: 5.3±2.3 SD). Proteins with a match were further analysed for secretion signals and eukaryotic-like domains (EffectorT3 model in EffectiveDB; Eichinger *et al*., 2016). Of the matched proteins, the EffectorT3 model supported 76 and 7 as effectors with high (>99%) or medium confidence (>95%), respectively. Some of the prominent domains that were confirmed to occur in effectors in various bacterial families were Shikimate kinase, DUF1512, glycosyl hydrolases and glycosyl transferases, NolW and NolX domains and metallo-beta-lactamase (Fig. S4).

The T5aSS, T5bSS and T5cSS were present in 166, 137 and 56 of the At_microbiome genomes, respectively, but the number of copies per genome varied greatly within and between families (Table S6). The highest T5aSS copy number was observed in the genomes of the *Bradyrhizobium* genus in the *Bradyrhizobiaceae* family (4-12 copies), for T5bSS in unclassified genera in the *Burkholderiaceae* (up to 9 copies) and for T5cSS in the *Variovorax* genus in the *Comamonadaceae* family (up to 8 copies).

The T5aSS and T5cSS are autotransporter proteins that code for a transporter domain (anchored in the membrane and facilitating translocation) and a passenger domain which is presented outside the membrane or secreted into the environment (Wilhelm *et al*., 2011). The most prominent T5aSS passenger domain identified in the genomes was an immunoglobulin (Ig) domain, which was generally present as tandem repeats. This domain has predominantly been associated with host cell attachment, when present on the outer membrane (Luo *et al*., 2000, Bodelón *et al*., 2013). Based on the predicted domains, T5aSS’s could be divided into different general functional categories, including host cell attachment, membrane hydrolysis, nuclease toxin and various adhesins. The functions differed between genera (Fig. S5A), for example *Variovorax* mostly contained extended signal peptides that contribute to inner membrane translocation (Leyton *et al*., 2012), *Cupriavidus* had some extended signal peptides but mostly domains for host cell attachment. *Brevudimonas* and *Sphingomonas* had subtilase and acylhydrolase domains, suggesting a role as toxins rather than host attachment in these genera.

The T5bSS is a two-partner system where the gene for the transmembrane protein is located next to the secreted protein gene, in the same operon on the genome (Fan *et al*., 2016). The predominant domains on potential exported proteins were Hemagglutinin domains and MGB/YDG domains, which were generally attached to the Hemagglutinin proteins (Fig. S5B). This protein forms part of the contact-dependent growth inhibition (CDI) system, playing a role in short-range interbacterial competition (Willett *et al*., 2015). Another prominent domain was a collagen middle domain, identified in genera of the *Burkholderiaceae* family including *Cupriavidus*. Collagen-like bacterial proteins have been linked to attachment and biofilm formation of *Burkholderia* species in animals (Bachert *et al*., 2015) and attachment to *A. thaliana* roots in a *Bacillus* species (Zhao *et al*., 2015).

T5cSS only contained extended signal peptides and adhesins domains, supporting that these systems solely contribute to cellular adhesion. This system was less prevalent in most of the At_microbiome genomes but had a high copy number in some of the *Burkholderia* (3-5 copies) and *Variovorax* isolates (max 8 copies).

The T6SSi was identified in 103 genomes. T6SS effectors can either be attached to the needle tip (cargo effectors) and are thus encoded as an additional domain of this gene in the genome, or will be secreted by the needle structure but are encoded as separate genes which can be found near the needle tip genes (*VgrG*/*PAAR* or *Hcp*) in the genome. A total of 107 cargo effectors were predicted in the genomes, ranging from one to seven copies per genome. Another 274 other transported effectors were identified, ranging from 1-14 effectors per genome (average: 3.3±3.0 SD). Based on functional domains, the functions predicted for the effectors (corresponding to those previously reported in literature; Table S7) ranged from various anti-bacterial toxins, carboxypeptidase, DNase activity, some eukaryotic targets, membrane pore formation, methyltransferase, peptidoglycan cleavage to post-translational modification (Fig. S6). Proteins with various different DUF domains, with unknown functions but previously associated with effector proteins (see Table S7), were also identified in the genomes. The most abundant effectors were those linked to peptidoglycan cleavage and antibacterial toxins.

### Multiple copies of the T6SSi can be placed in different, known, phylogenetic clusters

Multiple copies (2 - 3) of the T6SSi in a single genome were specifically identified in the *Burkholderiaceae, Oxalobacteraceae* and *Comamonadaceae* families (Fig. 5; Table S8). The T6SS has been subdivided into five known phylogenetic clusters which can be visualised in phylogenetic trees (Durán *et al*., 2021). Groups 1, 3 and 4 are commonly found in plant-associated microbes while some others might be predominantly animal-associated. To determine the classification of the different copies of T6SSi’s found here, we constructed a phylogenetic tree from all the translated *tssB* genes found in complete T6SSi clusters in the 241 At_microbiome genomes. The T6SS’s predominantly grouped into the Group 2, 3 and 4A clusters (Fig. 5).

**Figure 5.**
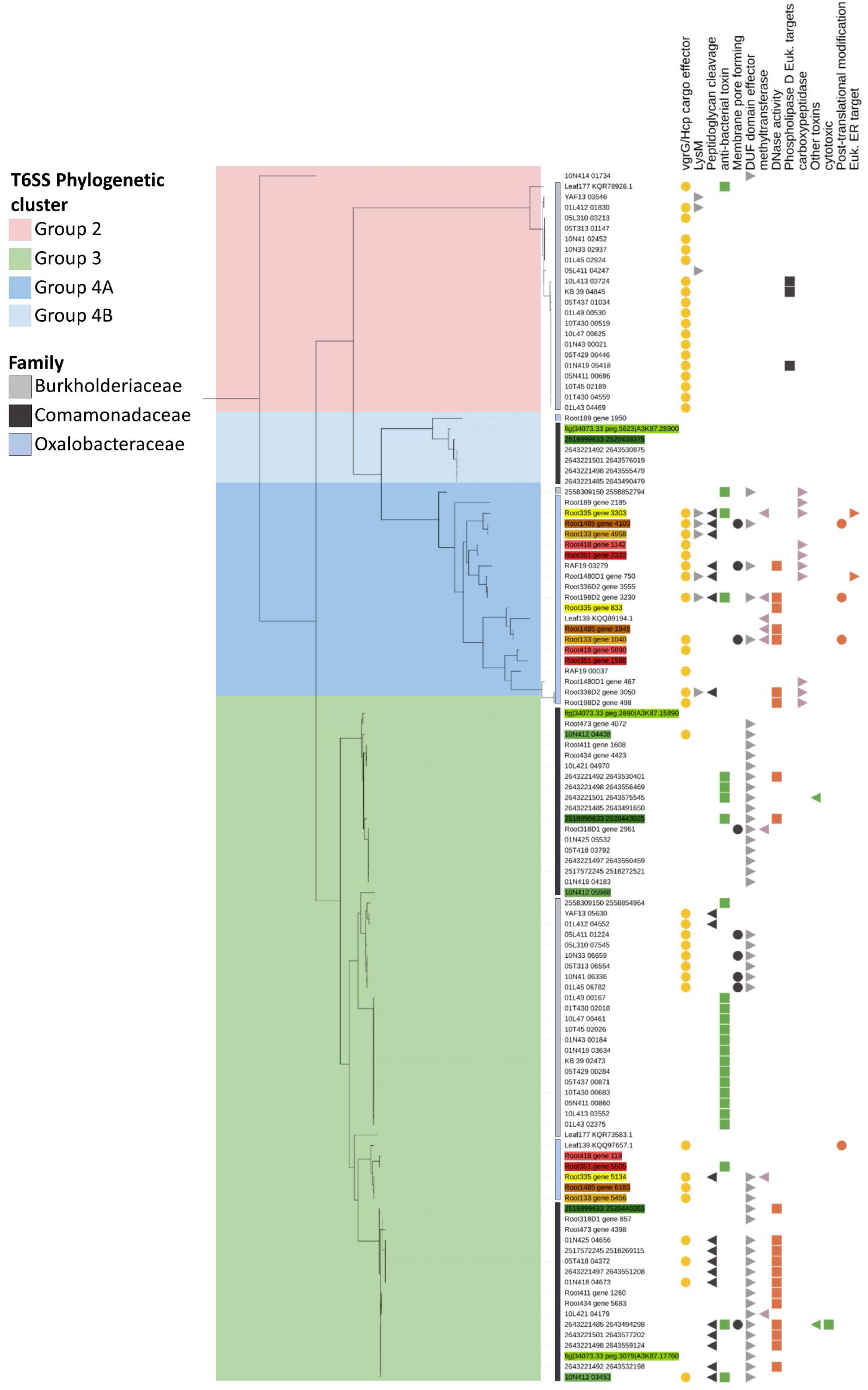
Summary of isolates encoding two or three T6SS copies in their genomes. Different effector functions can be linked to the different copies of T6SS’s. The ML tree was constructed based on the *tssB* gene. Coloured ranges on the branches indicate the well-known T6SS taxonomic groups. Coloured strips before the labels (gray scales) indicate taxonomic identity. Isolates with three T6SS copies are highlighted with coloured labels, each colour corresponding to a single genome. The different effector functions are summarised by the different coloured shapes.

In genomes that had multiple copies of the T6SSi, indicated with coloured labels in Fig. 5, the copies were often in different phylogenetic groups, suggesting complementary functions. For example, two of the copies were in subgroup 3 while the third was in subgroup 4A (*Oxalobacteraceae*) or in 4B (*Comamonadaceae*). This could indicate independent origins of the different copies. Indeed, the effector proteins associated with the different copies also had different functions (Fig. 5). One copy in *Oxalobacteraceae* genomes, for example, was associated with DNase activity, methyltransferase and post-translational modifications, while the second copy only contained *VgrG*/*Hcp* cargo effectors and proteins linked to peptidoglycan cleavage and the third copy only encoded a V*grG*/*Hcp* cargo effector and a protein with a DUF domain previously linked to T6SS effectors.

### Four SS’s are potentially active in the rhizosphere, based on *A. thaliana* metatranscriptomic data

Metatranscriptomic data were available for the same samples used in the *A. thaliana* metagenomic study (Sanchez-Gil *et al.,* Unpublished). These data were used to determine which SS’s were expressed in the soil or rhizosphere. Read mappings were converted to TPM (see M&M). The SS’s had a relatively low TPM value compared to those of four housekeeping genes, but active SS’s were clear for T1SS, T3SS, T5aSS and T9SS (Fig. S7), with the highest TPM values for T3SS and T5aSS. Expression was detected for eight of the nine core T3SS genes and six of these were significantly enriched (FDR <0.05) in the rhizosphere relative to the soil (logfold 0.7 – 1.8). For the T9SS, seven of the eight core components were expressed and all were more abundant in the rhizosphere (logfold 1.6 – 4.1). Two T6SSi genes, *tssD* and *tssB,* had very high expression levels (Fig. S7). The *tssD* (*Hcp*) gene often occurs independently from the gene cluster, in close proximity to secreted effector proteins or fused to effector proteins and can occur as multiple copies in a genome. This is one explanation for the high expression values. Nonetheless, this gene is considered as a marker to indicate if a T6SS is functional (Bernal *et al*., 2017), suggesting the high levels detected in this study relate to active T6SS’s in the rhizosphere.

## Discussion

Various bacterial SS’s, that are enriched and thus likely contribute to survival in the rhizosphere, were identified in this study. Although several studies have investigated SS’s in the genomes of plant-associated bacteria, few have considered their presence in natural root communities (microbiomes) of different plant species (Ofek-Lalzar *et al*., 2014, Levy *et al*., 2018). In this multi-plant, comparative metagenomic study, we found that the T2SS, T3SS, T5SS, T6SS and T9SS were more abundant in the rhizosphere community than in bulk soil in at least one plant species. The bacteria encoding these SS’s are more abundant near the plant root and likely benefit from the secreted effectors. Enrichment was family-specific and a number of bacterial families were identified that make use of these systems. The potential functions of the SS’s were investigated in species of interest, based on the predicted effector proteins, and some prominent functions included a role in interbacterial competition and host cell attachment.

Five of the SS types were more abundant in the rhizosphere relative to the bulk soil. Two of these systems, T3SS and T6SS, have received most attention in plant-associated microbiome studies as potential contributors to rhizosphere competence (Bernal *et al*., 2018, Borrero de Acuña & Bernal, 2021, Durán *et al*., 2021, Zboralski *et al*., 2022). The current study supports the importance of both these SS’s, as they were enriched in three to four different metagenomic datasets. T3SS’s also had higher expression levels in the *A. thaliana* rhizosphere than in bulk soil.

The collection of effector proteins predicted for T3SS and T6SS paints a picture of the various activities likely occurring simultaneously in such a community. Most predicted T6SS effectors had a toxic effect towards other bacteria that can result in cell lysis or DNA damage. Various *Cupriavidus* species have biocontrol activity against fungi (Ye *et al*., 2020, Estoppey *et al*., 2022) and indeed we observed *Cupriavidus* effectors with eukaryotic targets. Functional domains were also identified in T3SS effectors, providing some indication of their role. Functions such as glycosyl hydrolase and glycosyl transferase have been suggested to facilitate plant-microbe interactions (such as root cell penetration) in some genera (Wang *et al*., 2022) and the NolX domain detected in many effectors can serve as a translocation channel that facilitates transportation of effectors into plant cells (Staehelin & Krishnan, 2015). The shikimate kinase domain, present in many of the effectors in the current study, has also been found in a T3SS effector of a *Bradyrhizobium* sp. (Piromyou *et al*., 2021) and a *Mesorhizobium* sp. (Okazaki *et al*., 2010). In the latter species one such effector triggered bacterial recognition in the host plant (*Lotus halophilus*) which limited bacterial colonisation and reduced nodulation. Nonetheless, this recognition suggests that this effector domain might also facilitate colonisation of other hosts.

Furthermore, additional SS’s were identified as important in the rhizosphere, namely T2SS, T5SS, *Bacteroidetes*-specific T6SSiii, and T9SS. T2SS’s has mostly been investigated in plant pathogens but also occurs in environmental microbes, facilitating the secretion of CAZymes for cell wall degradation, lipases and proteases that can facilitate plant host colonisation (Cianciotto & White, 2017, Pfeilmeier *et al*., 2023, Entila *et al*., 2024). The different T5SS’s have rarely been investigated in plant systems but could be big contributors to microbial establishment on the host. In this study, the most abundant functional domain of T5aSS was the Ig domain, associated with host cell attachment (Luo *et al*., 2000, Bodelón *et al*., 2013). The T5aSS is well known for its contribution to biofilm formation and adhesion to plant surfaces in some plant-associated microbes (Castiblanco & Sundin, 2016). Its functional importance has been confirmed in *Xanthomonas* pathogens (Alvarez-Martinez *et al*., 2021) but it has not been studied in other genera. The Hemagglutinin domain was prominent in the T5bSS, a known signature of the contact-dependent growth inhibition (CDI) system. The T5bSS likely also contributes to interbacterial competition or host interaction, as observed in animal hosts (Allen *et al*., 2020), but this still requires thorough investigation in plant rhizospheres.

The rhizosphere-enriched T6SSiii and T9SS were predominantly assigned to members of the *Chitinophagaceae* family, for which the role of their SS’s in plant interactions has rarely been investigated. This family is also considered as part of the core microbiome in the rhizosphere and endosphere of different plant species (Schneijderberg *et al*., 2020). The transcriptomic data from *A. thaliana* suggested active expression of the T9SS in the rhizosphere, supporting the involvement of this SS in rhizosphere interactions. T6SSiii plays a similar role as other T6SS’s, i.e. producing a needle-like injection structure to attack nearby microbes (Russell *et al*., 2014), while the T9SS can contribute to gliding motility (i.e. adhesins) and secretion of virulence factors and chitinases (Song *et al*., 2022). *Chitinophagaceae* genera can be beneficial to plants directly, or serve as biocontrol agents against fungal pathogens (Madhaiyan *et al*., 2015, Carrión *et al*., 2019).

The enrichment of SS’s in the rhizosphere or soil could be linked to various specific bacterial families. Many of these are considered core taxa of plant microbiomes, such as *Rhizobiales, Burkholderiales, Pseudomonadales*, the *Bacteroidetes* phylum (Banerjee *et al*., 2018), *Oxalobacteraceae, Comamonadaceae* and *Xanthomonadaceae* (Schneijderberg *et al*., 2020, Johnston-Monje *et al*., 2021). A predominant focus in studies of T3SS and plant-associated bacteria has been on *Pseudomonas*, root-nodulating *Rhizobiales* and other genera within this order (Stringlis *et al*., 2019, Wallner *et al*., 2021, Boak *et al*., 2022, Zboralski *et al*., 2022). This study identified additional families likely benefiting from T3SS and T6SS, such as *Oxalobacteraceae, Comamonadaceae* and *Caulobacteraceae*. Various genera in these families are considered beneficial microbes (Wen *et al*., 2020, Berrios, 2021, Yu *et al*., 2021) and could be good candidates for the development of plant growth-promoting microbes or biocontrol agents.

The bacterial isolates encoding multiple copies of some SS’s were of specific interest, since these copies likely serve complementary functions and provide a larger arsenal of SS tools. This could specifically be investigated in the T6SS, since its corresponding effector genes are located in close proximity to the structural SS genes in the genome. In the *Burkholderiaceae, Oxalobacteraceae* and *Comamonadaceae* families, two to three copies were prominent in many isolates. The different copies often grouped in different phylogenetic groups and were linked to different secreted products and functions. Multiple copies have been shown to play different roles in plant growth-promoting microbes. For example, in *Pseudomonas chlororaphis,* two T6SS’s contribute to interbacterial competition and protection against predation but the two copies were controlled uniquely and had different bacterial target ranges (Boak *et al*., 2022). Similarly, in *Pseudomonas putida* three T6SS’s were found which collectively contribute to interbacterial competition but one copy was significantly more important for killing a broad range of bacteria than the other two (Bernal *et al*., 2017).

Enriched SS’s could not be identified robustly across all the studied datasets. In the *A. thaliana*, cucumber (Ofek-Lalzar *et al*., 2014) and wheat (Zhou *et al*., 2020) datasets, various enrichments were observed and the two studies of *A. thaliana* were quite consistent. An explanation for unclear distinctions in the other datasets could be due to the sampling method. In the former studies, the root and tightly adhering bacteria were included whereas in most of the latter studies the root-adhering soil was collected and the root excluded. This can result in smaller differences between the rhizosphere and bulk soil communities. This was also prominent in the alpha diversity differences between rhizosphere and soil communities, where the studies including the root in the sampling displayed a clearer rhizosphere effect (i.e., reduced alpha diversity in the rhizosphere). We thus suggest that alpha and beta diversity measurements should be analysed first and/or used for proper interpretation, as key indicators of the rhizosphere effect.

Our comparison of bacterial communities from paired rhizosphere and soil metagenomic samples of different plant hosts revealed an enrichment of many SS’s in the rhizosphere, and their functional interpretation highlighted several potential mechanisms by which rhizosphere bacteria might colonize and persist in the rhizosphere. Thus, our study provides a guide of which bacterial families can establish in the rhizosphere, and their SS’s that may contribute to this process. Essential functions such as interbacterial competition (T5bSS and T6SS) and attachment to the host surface (T5SS) seem to be important for improving establishment near the root. These SS’s can also be transferred to related species of importance to enhance their survival in the rhizosphere environment (Borrero de Acuña & Bernal, 2021). Families with these useful machineries can be mined for additional properties such as plant growth promotion or biocontrol, knowing they have survival benefits in a natural community.

## Materials and methods

### Metagenomic data processing and taxonomic composition

Metagenomic sequence data were gathered from publicly available datasets, including paired rhizosphere and soil collections (Ofek-Lalzar *et al*., 2014, Sanchez-Gil *et al.,* Unpublished, Stringlis *et al*., 2018, Xu *et al*., 2018, Zhou *et al*., 2020). Datasets with high quality paired-end Illumina sequence data and at least 30,000 reads per sample were selected for appropriate comparisons between studies. The rhizosphere and soil samples consisted, on average, of three biological replicates. Additional details on the datasets are provided in Table S1.

Metagenomic reads were processed, assembled and annotated using the ATLAS pipeline (Kieser *et al*., 2020). Quality filtering included read filtering and trimming based on sequence quality and the removal of host DNA, using appropriate host reference genomes (Table S1). Due to the size of the wheat reference genome an alternative approach was used to remove plant DNA from these datasets, using Kraken 2 (Wood *et al*., 2019), and only reads that did not match to the eukaryotic database were retained. Metagenome assembly was performed by metaSPAdes (Nurk *et al*., 2017) and genes were predicted with Prodigal (Hyatt *et al*., 2010), both included as part of ATLAS. To evaluate the coverage and assembly quality of the metagenomes, the additional step in the ATLAS pipeline was included, where the raw reads were mapped back to the assembly.

Taxonomic composition of the metagenomic datasets was determined by read classification, using Kaiju v. 1.6.2 (Menzel *et al*., 2016). The relative abundances of different genera and families were determined based on all classified bacterial reads. The relative abundances at genus rank were used for calculation of the alpha and beta diversity of the datasets. The Shannon diversity index was calculated as an indication of alpha diversity and differences in alpha diversity between datasets were tested with pairwise t-tests (FDR adjustment Benjamini & Hochberg). For comparison of beta diversity, Bray-Curtis dissimilarity was calculated and differences between datasets were tested with a permanova test, followed by pairwise comparisons with pairwiseAdonis (BH adjustments for FDR). For this analysis genera with relative abundance >1·10^-5^ were used. Dataset-specific comparisons between rhizosphere and soil samples of a dataset were also performed using the same approach. Packages used with R 4.0.3 (R Core Team, 2020) in RStudio (RStudio Team, 2019) included phyloseq (McMurdie & Holmes, 2013), metagMisc (Mikryukov, 2019), vegan v. 2.5-7 (Oksanen *et al*., 2013) and pairwiseAdonis (Arbizu, 2019).

### Identification of secretion systems in the metagenomic data

A read mapping approach was used to identify the SS’s in the metagenomes. To create the SS protein database for each metagenomic study, SS proteins were predicted in the metagenomic contigs as well as in the 254,090 GTDB genomes (release 202) (Parks *et al*., 2021). SS proteins were predicted with MacSyFinder (Abby *et al*., 2016), which uses an hmm search against the TXSScan models, including the T1SS - T6SS and T9SS and the subtypes T4SSI, T4SST, T5aSS, T5bSS, T5cSS, T6SSi, T6SSii and T6SSiii. In the GTDB genomes, complete SS’s were predicted using the gembase mode and in the metagenomes the unordered mode was used. The resulting collection of SS proteins was reduced to a non-redundant set, based on 98% similarity, using CD-hit v. 4.8.1 (Fu *et al*., 2012). Due to the ambiguity of some components of the SS’s (see Introduction), a subset of core genes was selected for many of the SS’s (see Table S3) and a DIAMOND BLAST database was generated for downstream analysis (Buchfink *et al*., 2015).

Quality-filtered reads of each metagenome sample were mapped to the SS protein database using the sensitive DIAMOND blastx algorithm (Buchfink *et al*., 2015) with >40% identity and e-value < 10^-5^ cut-offs. Results were processed in R Studio to only select genes with >60% coverage and converted to a normalised count, termed metagenomic reads per kilobase million (MRPM). This was based on the formula used for calculating transcripts per kilobase million (TPM), which first normalises for gene length and then for read depth. The term MRPM is used throughout this manuscript to avoid confusion with RNA transcripts. A kingdom-wide study of bacterial SS’s indicated that some SS’s only occur in specific phyla, such as the T2SS only in *Proteobacteria*, and T6SSiii and T9SS only in Bacteroidetes (Abby *et al*., 2016). Thus, the MRPM counts were adjusted based on the relative abundance of the phyla known to encode for such SS’s (using the formula (MRPM count)/(Phylum relative abundance)). Kaiju analyses of the reads, previously described, were used to determine the relative abundances of the phyla. The MRPM values of all orthologs of each SS gene were summed and the median MRPM value of each SS was determined from all its gene components.

For each plant metagenomic dataset, pairwise t-tests were performed to determine which SS’s were more abundant in the rhizosphere or the soil biome. Analyses were done in RStudio (RStudio Team, 2019), using the rstatix package and *p values* were corrected for FDR (Benjamini-Hochberg).

### Taxonomic classification of the identified secretion systems

The SS proteins identified in the metagenomes were classified up to family rank. The SS proteins from the assembled metagenomic contigs were annotated based on CAT classification of the contigs (using NCBI database) (von Meijenfeldt *et al*., 2019) and those identified on GTDB genomes were annotated based on the GTDB metadata (using NCBI taxonomic names for conformity). The total MRPM value of each SS protein was summed per taxonomic family. To determine the abundance of a SS in a given family, we used an approach similar to pathway predictions in the HUMAnN2 software (Franzosa *et al*., 2018). First, we replaced the lowest MRPM value of a SS protein with that of the second lowest value, as a gap filling approach to compensate for missing proteins or miss-annotation. Second, we calculated the abundance of a SS by averaging the values of all SS proteins in the family (HUMAnN2 used average of the top 50% reactions of a pathway). In order to focus mostly on complete SS’s in the families, a SS was only included if more than 40% of the proteins were detected for a specific family.

Enrichment of the SS’s of different families in the rhizosphere or soil were determined with DESeq2 v.1.28.1 (Love *et al*., 2014), using the average calculated MRPM values. A size factor of 1 was used to bypass the normalisation step, since MRPM values were already normalised. Families with SS’s that were enriched in the rhizosphere or soil biome in multiple plant species were considered as the families most likely benefitting from the SS’s.

### Functional prediction of SS’s in prominent bacterial families - *Arabidopsis thaliana* as model

More detailed investigations of the secreted protein products were performed on publicly available *A. thaliana* isolated genomes (At_microbiome genomes). Genome sources used in this study included IMG (JGI Integrated Microbial Genomes Database; Chen *et al*., 2022), PATRIC (BV-BRC) database version 2022-02 (Wattam *et al*., 2017), the At-Sphere collection from the MPIPZ institute (Bai *et al*., 2015) and a local culture collection from Utrecht University, constructed from limed Reijerscamp soil (Selten *et al*., 2024). Families with SS’s that were enriched in the bulk soil or rhizosphere biomes (T2SS, T3SS, T5SS and T6SS) were selected for further analyses, including 241 genomes (Table S4) from the families: *Rhizobiaceae, Comamonadaceae, Burkholderiaceae, Caulobacteraceae, Sphingomonadaceae, Bradyrhizobiaceae* and *Oxalobacteraceae*.

### Prediction of secreted proteins of T3SS, T5SS and T6SS

Secreted proteins were predicted from the previously described At_microbiome genomes resource (Bai *et al*., 2015, Wattam *et al*., 2017, Chen *et al*., 2022, Selten et al., 2024). Functions of putative secreted proteins were predicted by comparison to the Pfam database of protein families (Mistry *et al*., 2020), using InterProScan v. 5 (Jones *et al*., 2014).

For the T3SS, effector genes are not necessarily located close to the T3SS structural gene cluster in a genome, therefore predictions were based on similarity to verified effector proteins (Hu *et al*., 2017). The protein sequences from all the genomes with a confirmed T3SS were compared to a previously generated effector database (Hu *et al*., 2017) using blastp with a >30% identity cutoff. The potential effector proteins were further analysed with the EffectiveT3 model in EffectiveDB (Eichinger *et al*., 2016) for effector signals such as secretion signals and eukaryotic-like domains.

T5aSS and T5cSS are autotransporters that encode for a transporter domain and a passenger domain (Wilhelm *et al*., 2011). The functional annotations of the passenger domains were used to predict the putative functions of the SS’s. The T5bSS is a two-partner system where the gene for the outer membrane protein is located next to the secreted protein gene in the same operon on the genome (Fan *et al*., 2016). Secreted proteins of T5bSS were predicted by collecting all proteins flanking the T5bSS outer membrane protein in the genome. Several functions and domains have been reported in literature for T5bSS (Table S7) (Barret *et al*., 2011), including contact-dependant growth inhibition (CDI) proteins (Willett *et al*., 2015, Lin *et al*., 2020), hemolysins, adhesin proteins, Ig domains (Belikov *et al*., 2021), collagen (Bachert *et al*., 2015) and tyrosine phosphatase (Rojas-Lopez *et al*., 2018). Proteins were filtered based on these predicted functions and potential novel functions.

The T6SS effectors are often attached to the needle spike protein VgrG or PAAR (C-terminal) and transported as a passenger on the needle tip. Thus the genes of the secreted effectors are either fused, or in close proximity to the *VgrG* or *Hcp* genes on the genome. Putative effectors were predicted by identifying 10 flanking genes on each side of *VgrG* and *Hcp* genes on the genomes and predicting their functional domains. Functional domains that have been associated with T6SS effectors in literature and associated with functions such as peptidoglycan cleavage, nuclease activity, membrane pore formation, interference with energy balance or post-translational modification or various known domains of unknown function (DUF) were used to identify potential effector candidates (Table S7) (Suarez *et al*., 2010, Salomon *et al*., 2014, Jiang *et al*., 2016, Klockgether & Tümmler, 2017, Verster *et al*., 2017, Bayer-Santos *et al*., 2019, Durán *et al*., 2021, Jurėnas & Journet, 2021, Boak *et al*., 2022). Predicted secreted proteins smaller than 50 amino acids were excluded from the analysis.

### Phylogenetic clusters and copy number of the T6SS’s

T6SS’s can be subdivided into known phylogenetic clusters, based on the phylogenetic composition of core structural genes. Five clusters have been well-defined in different studies (Group 1-5)(Bernal *et al*., 2018, Durán *et al*., 2021). To obtain the overall phylogenetic composition of this SS in the At_microbiome genomes, the most frequently used phylogenetic marker gene was selected for tree construction (*tssB*). Reference sequences from previous publications were included to easily identify known clusters. Amino acid sequences of the proteins were aligned using MAFFT v 7.453 (Katoh & Standley, 2013), large, gapped regions were removed with trimAl v1.4 (Capella-Gutiérrez *et al*., 2009), using the heuristic automated setting, and maximum likelihood trees were constructed with IQ-TREE v. 2.1.4 (Minh *et al*., 2020), including 1000 replicates of ultrafast bootstrap approximations. Trees were visualised and edited in iTOL v.5 (Letunic & Bork, 2021). The number of T6SS’s identified in a single genome was also investigated to identify which isolates and genera encode for multiple copies of a T6SS.

### Identification of “active” secretion systems in *A. thaliana* metatranscriptomic data

The metatranscriptomic data, from the same samples used in the *A. thaliana* metagenomic study (Sanchez-Gil et al., Unpublished), were used to investigate SS expression. Reads were filtered by removing rRNA sequences using SortMeRNA v. 4.3.4 (Kopylova *et al*., 2012) and host reads were removed using Kraken 2 (Wood *et al*., 2019), as described earlier for the metagenomic datasets. Transcriptome assembly was performed with Trinity v. 2.13.2 (Grabherr *et al*., 2011) with a minimum contig length of 150 bp, a maximum cluster size of 25 was set for the Chrysalis contig clustering step and reads were not normalised. Prediction of coding regions in the assembled transcripts were performed with TransDecoder v. 5.5.0 (Haas, 2023). The second step of CDS prediction with TransDecoder included information on CDSs with hits to the Pfam and Swissprot database as a prior for selecting the best CDSs. A single best CDS was retained per transcript.

The identified CDSs were reduced to a non-redundant set (nrCDS) at a 95% similarity threshold using CD-Hit-est (Fu *et al*., 2012). Reads were mapped to the nrCDS sequences using Bowtie2 (Langmead & Salzberg, 2012) and mapped reads were counted using HtSeq v. 0.11.3 (Putri *et al*., 2022) with the parameters --nonunique all and --mode union. Read counts were normalised by conversion to transcripts per kilobase million (TPM). All CDSs assigned to the same SS protein were combined for a total count per SS protein and TPM values were compared between the rhizosphere and soil biome samples. Four housekeeping genes (*atpD*, *ftsZ*, *rho* and *rpoA*) were included as a reference indication of the expressions levels that can be obtained for a gene in metatranscriptomic data. Differential expression of each SS protein in the rhizosphere versus the soil were compared with DESeq2 v.1.28.1 (Love *et al*., 2014), using the raw read counts as input.

## Data and scripts availability

The scripts used in this study can be accessed at the GitHub repository: https://github.com/Arfourie/Bacterial_SS's. Sequence data from the *A. thaliana* metagenome and metatranscriptome study by Sanchez-Gil *et al*. (Unpublished) will be made available at NCBI SRA upon publication. Genome sequences of isolates from the *A. thaliana* rhizosphere from the UU repository can be accessed at Zenodo (Selten *et al*., 2024).

## Acknowledgements

This study was supported by the Netherlands Organization for Scientific Research (NWO) Groen II project number ALWGR.2017.002, MiCRop Consortium programme, Harnessing the second genome of plants (Grant number 024.004.014), the European Research Council (ERC) Consolidator grant 865694: DiversiPHI, the Deutsche Forschungsgemeinschaft (DFG, German Research Foundation) under Germany’s Excellence Strategy – EXC 2051 – Project-ID 390713860, and the Alexander von Humboldt Foundation in the context of an Alexander von Humboldt-Professorship founded by German Federal Ministry of Education and Research.

---

**Figure S1.**
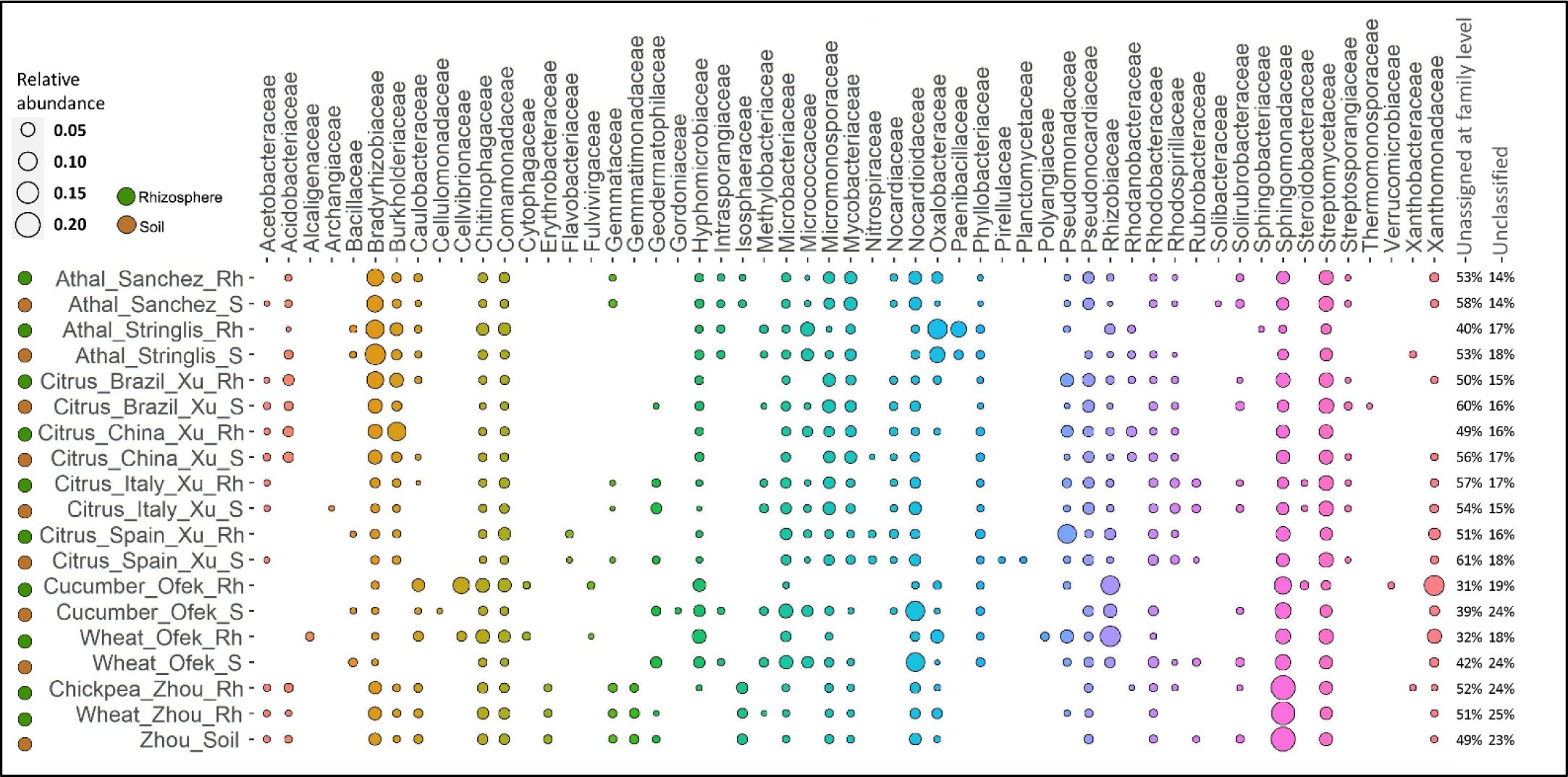
Families with > 1% relative abundance in the classified bacterial reads of the metagenomes of the different plant species (Kaiju read classification). Some plants had clear distinctions between rhizosphere (Rh) and soil (S), while others were highly similar in both biomes. Some of the most abundant families in all plants were *Bradyrhizobiaceae, Burkholderiaceae, Sphingomonadaceae* and *Streptomycetaceae*.

**Figure S2.**
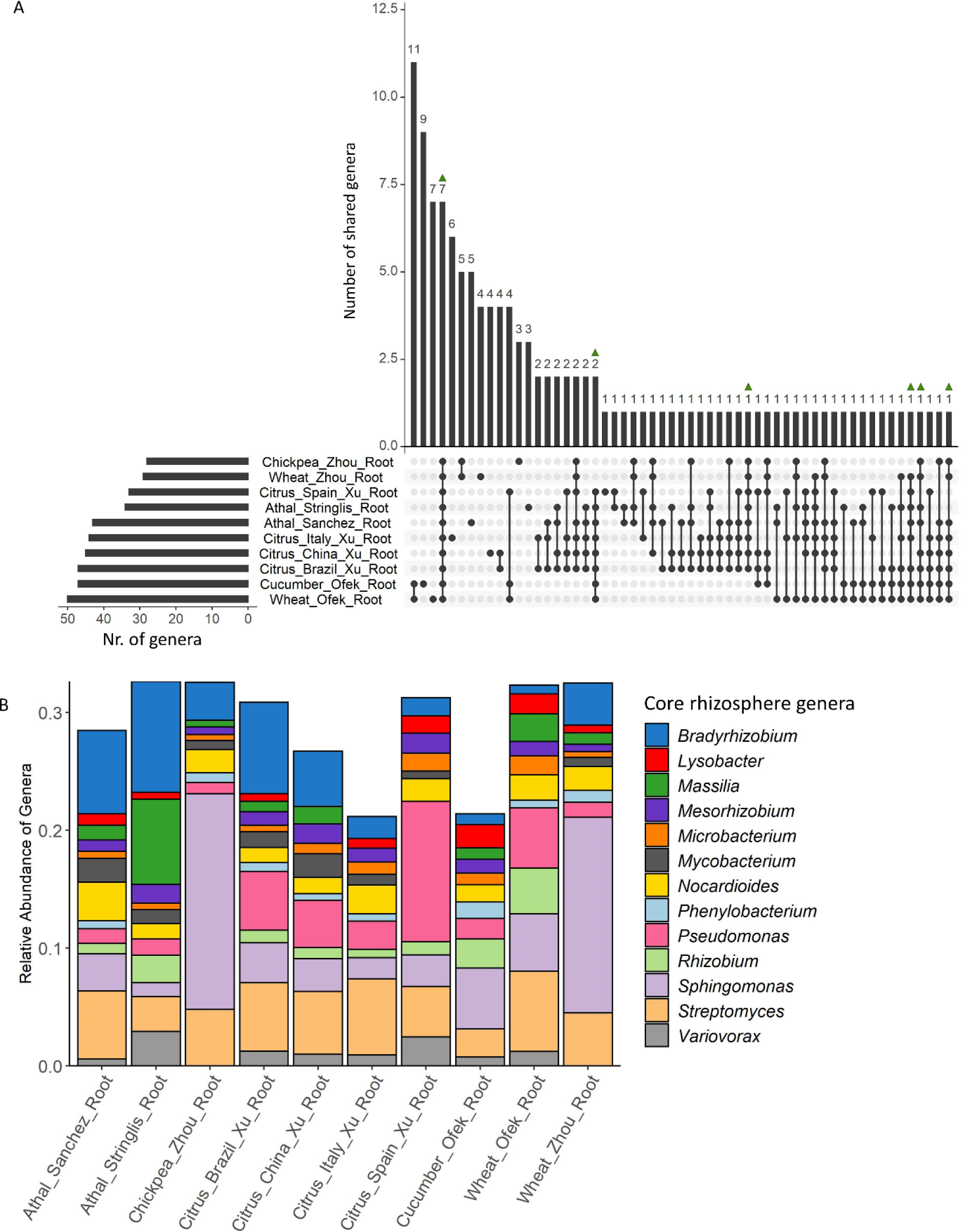
Summary of core bacterial genera (> 0.5% relative abundance) in the rhizosphere of all plant datasets, based on all bacterial reads classified at genus rank (Kaiju). A) The number of core genera and genera unique to each study is shown by connected dots. Green triangles indicate the genera present in >80% of all rhizosphere datasets (core genera) B) The relative abundance of the 13 core genera in the rhizospheres of the different datasets are illustrated in the coloured bar graph.

**Figure S3.**
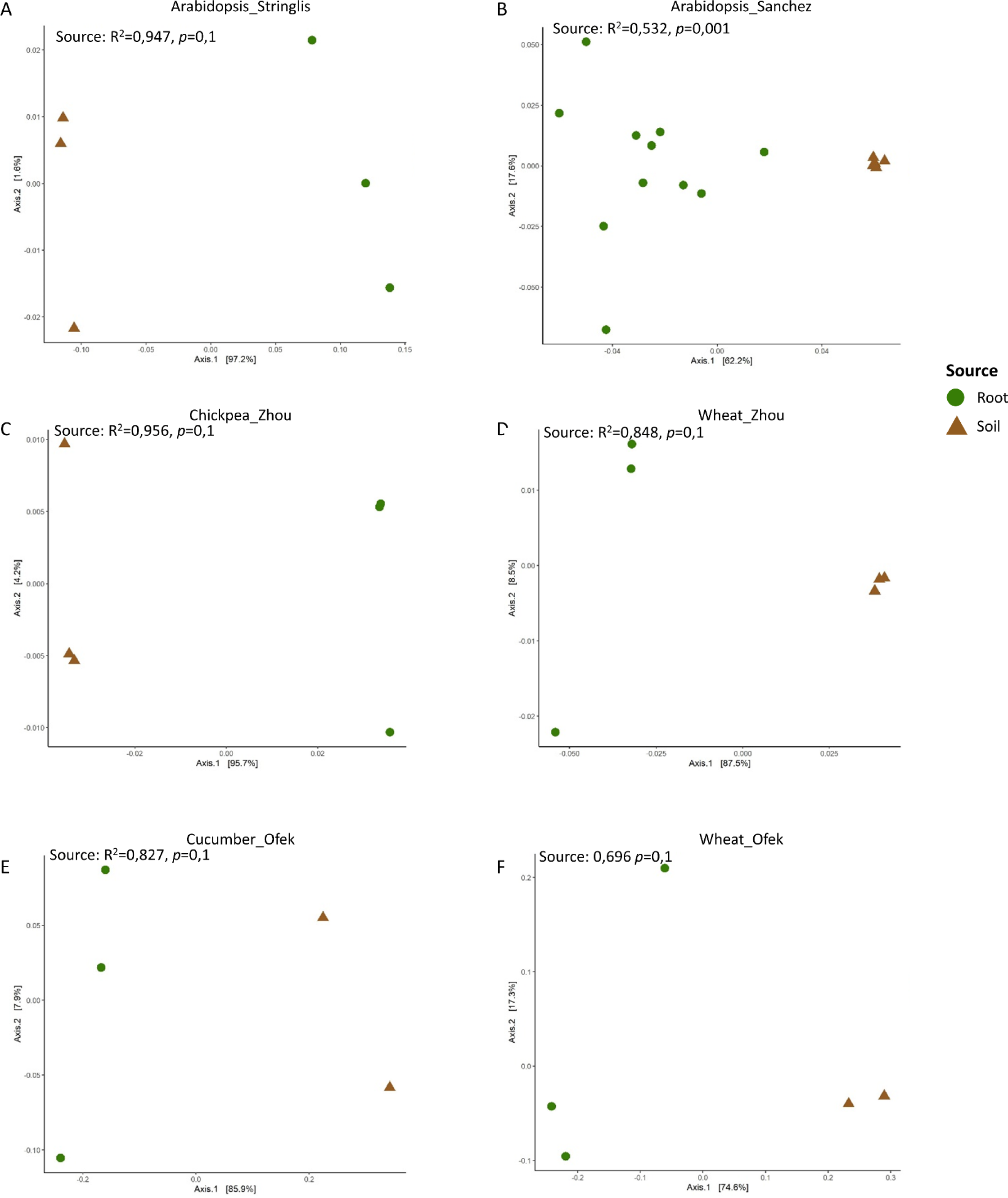
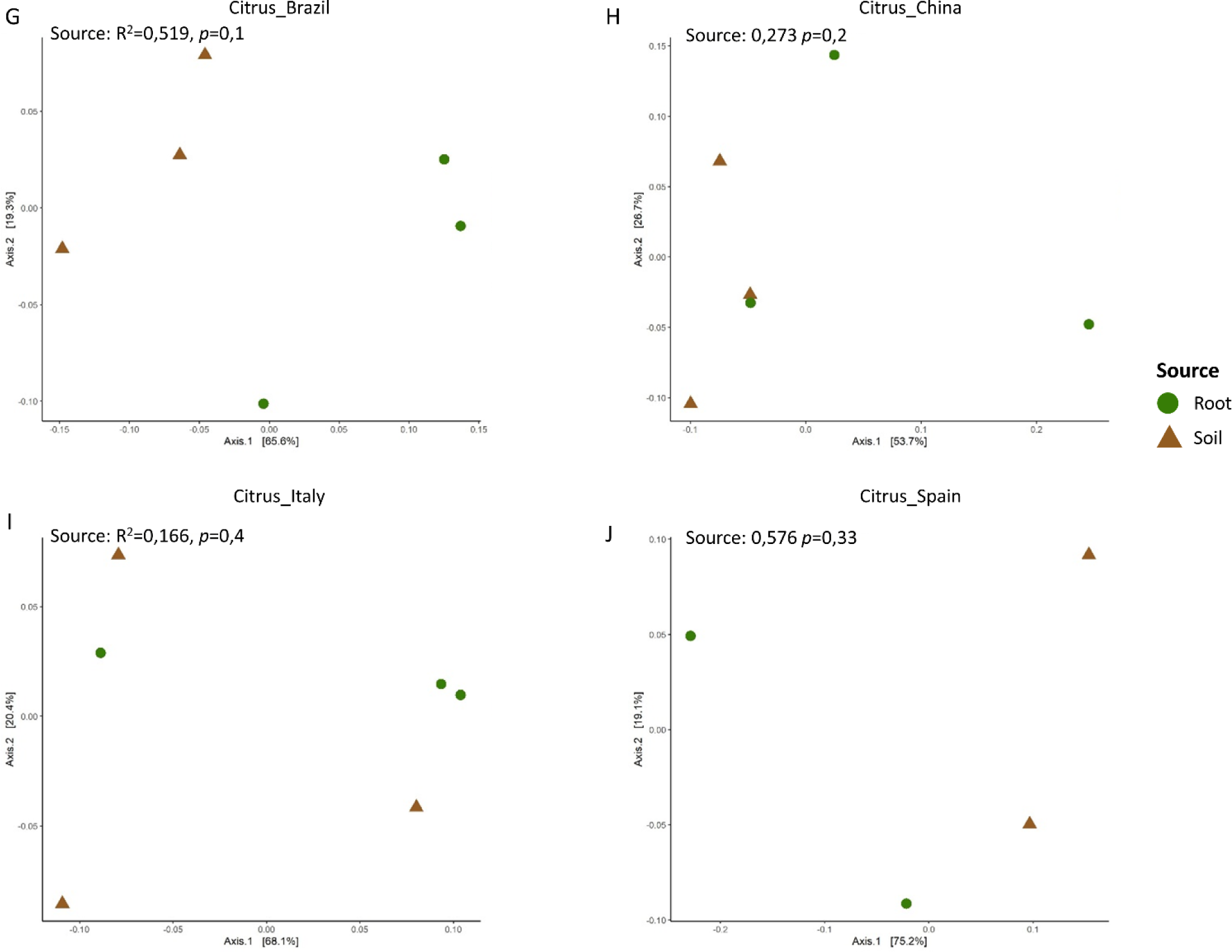
Dataset-specific beta-diversity analyses. The Bray-Curtis distances between samples were used to construct a principal coordinate analysis plot (based on genera > 1e-5 relative abundance) to illustrate differences in community composition between rhizosphere and soil in each dataset, including A) Arabidopsis_Stringlis, B) Arabidopsis_Sanchez, C) Chickpea_Zhou, D) Wheat_Zhou, E) Cucumber_Ofek, F) Wheat_Ofek, G) Citrus_Brazil, H) Citrus_China, I) Citrus_Italy, J) Citrus_Spain

**Figure S4.**
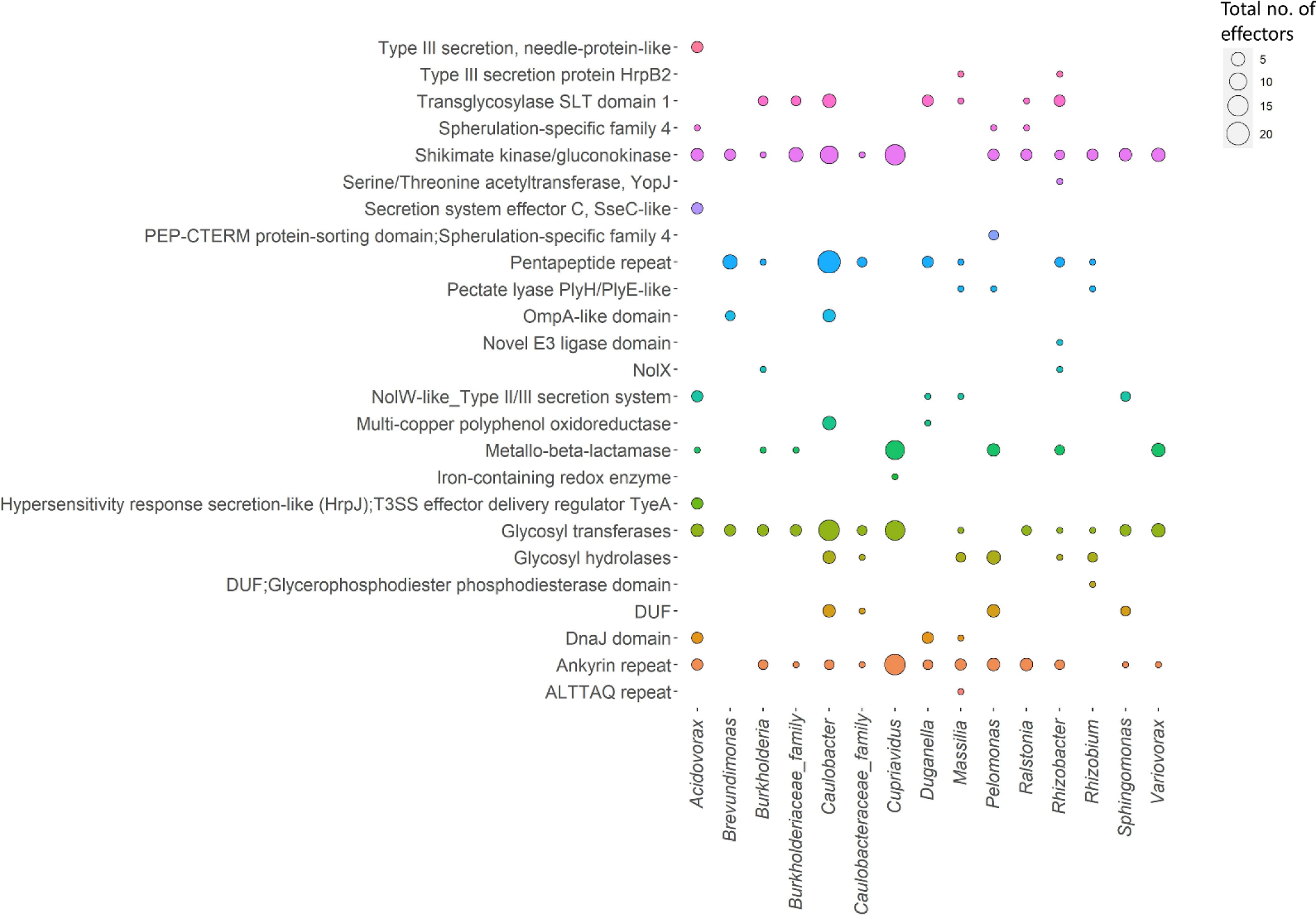
Summary of the functional T3SS effector domains present in different genera of the *A. thaliana* isolated bacterial genomes. Dot sizes represent the number of effectors predicted in the genus and the colours correspond to the functional domains.

**Figure S5.**
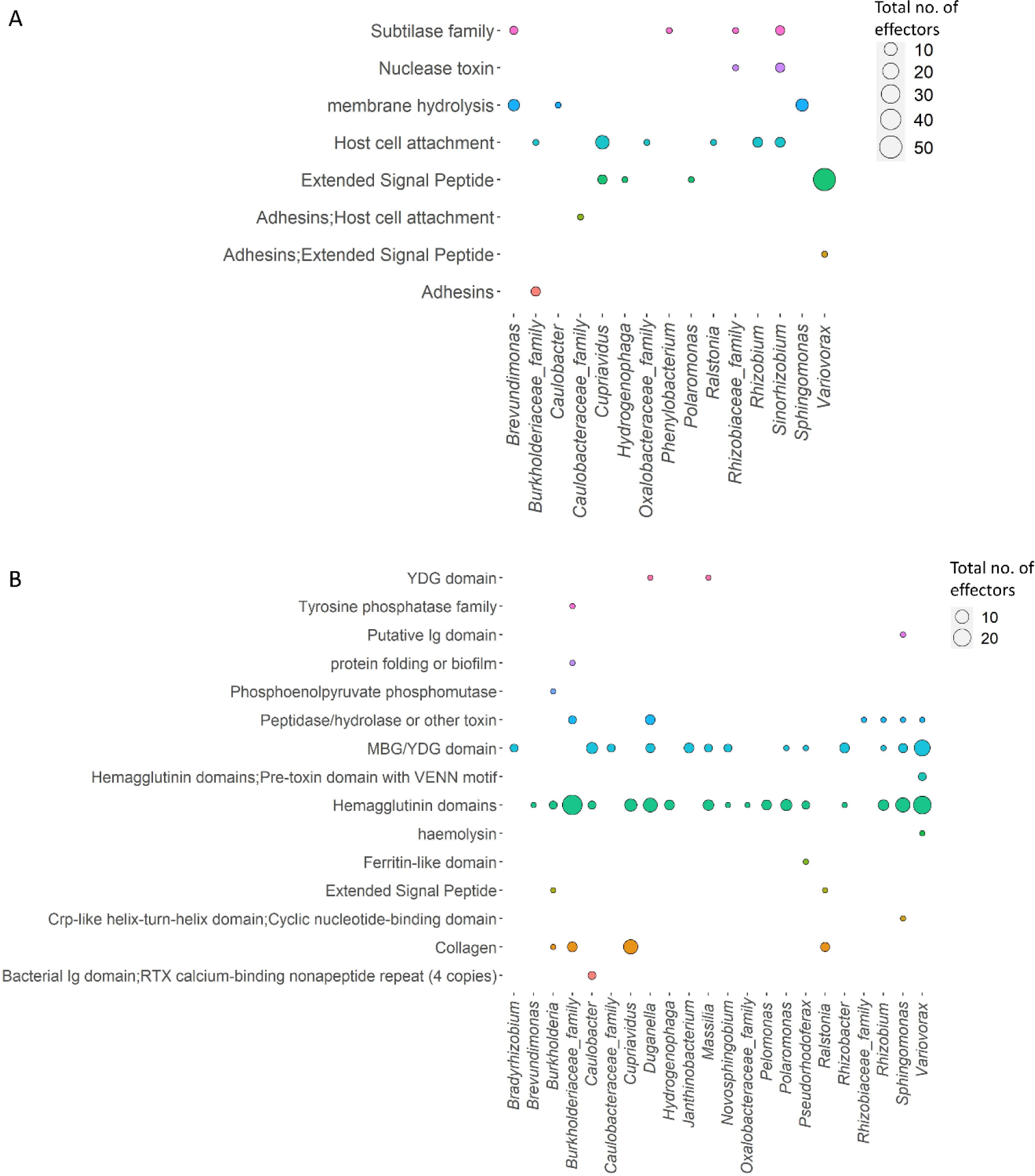
Summary of the functional domains/categories identified for the T5SS effectors, indicating the abundance of the categories in the (A) T5aSS and (B) T5bSS. Dot sizes represent the number of effectors predicted in the genus and the colours correspond to the functional domains.

**Figure S6.**
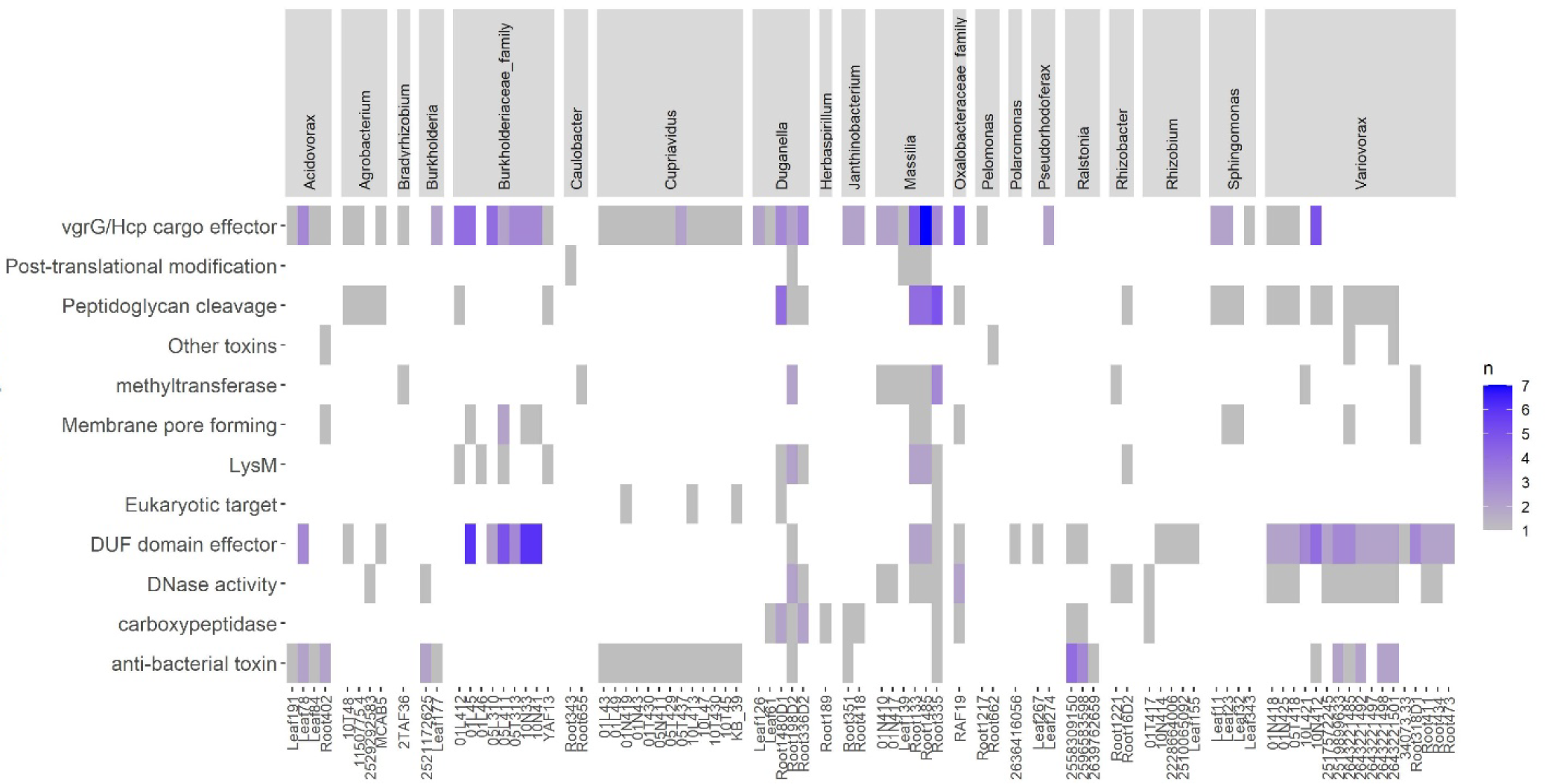
Summary of the effector functions identified in families with a T6SS and enriched in the rhizosphere.

**Figure S7.**
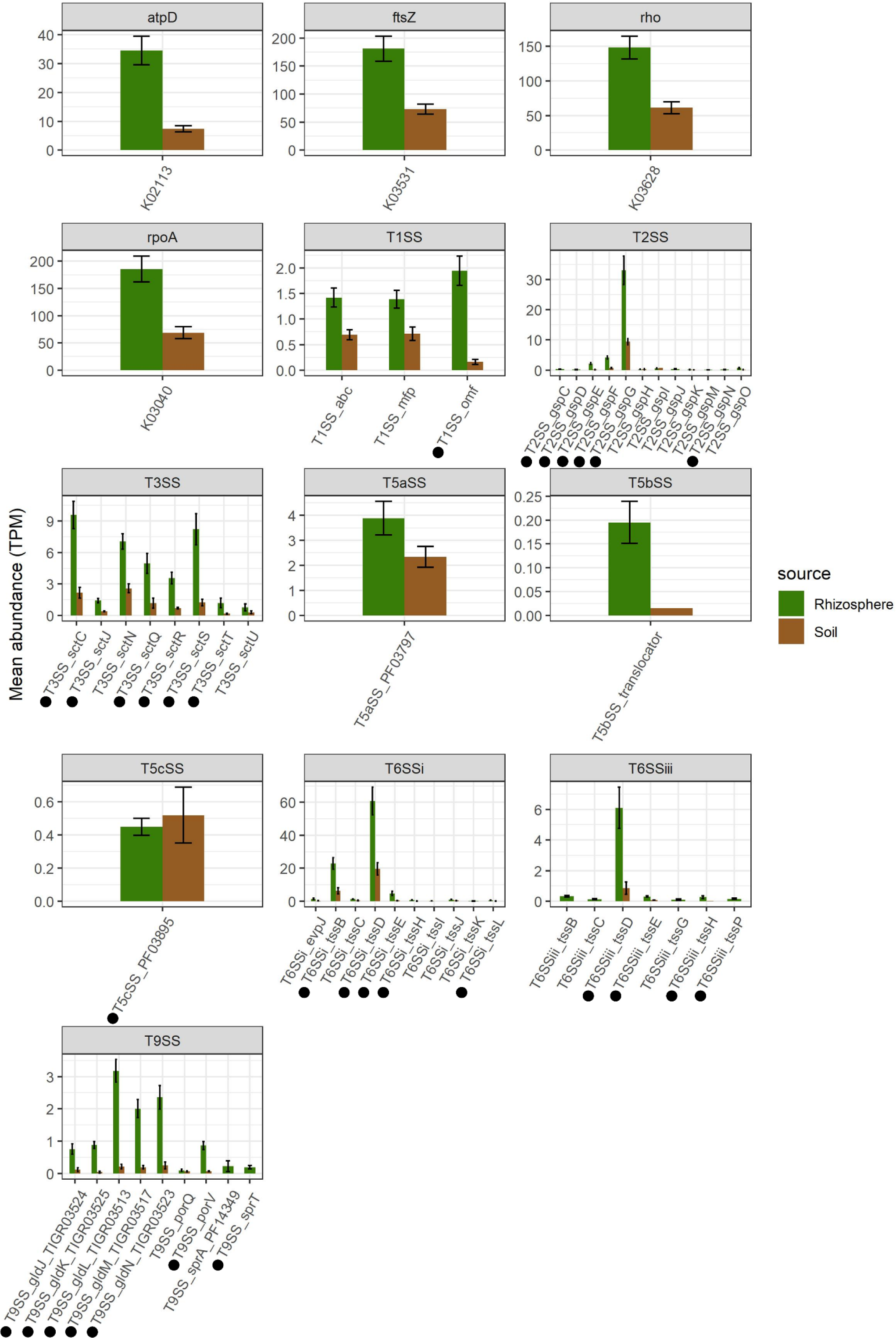
Expression of all the SS’s (and four housekeeping reference genes) in the metatranscriptomic dataset of *A. thaliana* rhizosphere and soil. Read counts were converted to TPM values to normalise reads between samples. The protein orthologs with significant logfold differences between the rhizosphere and soil samples (DESeq2, FDR <0.05) are indicated with a black dot.

## References

Abby SS, Cury J, Guglielmini J, Néron B, Touchon M & Rocha EPC (2016) Identification of protein secretion systems in bacterial genomes. Scientific Reports 6: 23080.

Abdallah AM, Gey van Pittius NC, DiGiuseppe Champion PA, Cox J, Luirink J, Vandenbroucke-Grauls CMJE, Appelmelk BJ & Bitter W (2007) Type VII secretion — mycobacteria show the way. Nature Reviews Microbiology 5: 883–891.

Allen JP, Ozer EA, Minasov G, Shuvalova L, Kiryukhina O, Anderson WF, Satchell KJF & Hauser AR (2020) A comparative genomics approach identifies contact-dependent growth inhibition as a virulence determinant. Proceedings of the National Academy of Sciences 117: 6811–6821.

Alvarez-Martinez CE, Sgro GG, Araujo GG, Paiva MRN, Matsuyama BY, Guzzo CR, Andrade MO & Farah CS (2021) Secrete or perish: The role of secretion systems in *Xanthomonas* biology. Computational and Structural Biotechnology Journal 19: 279–302.

An Y, Wang J, Li C, Leier A, Marquez-Lago T, Wilksch J, Zhang Y, Webb GI, Song J & Lithgow T (2016) Comprehensive assessment and performance improvement of effector protein predictors for bacterial secretion systems III, IV and VI. Briefings in Bioinformatics 19: 148–161.

Arbizu PM (2019) pairwiseAdonis: Pairwise multilevel comparison using adonis R package version 0.3. *See* https://github.com/pmartinezarbizu/pairwiseAdonis.

Ates LS, Houben ENG & Bitter W (2016) Type VII Secretion: A highly versatile secretion system. Microbiology Spectrum 4: 4.1.12.

Bachert BA, Choi SJ, Snyder AK, et al. (2015) A unique set of the *Burkholderia* collagen-like proteins provides insight into pathogenesis, genome evolution and niche adaptation, and infection detection. PLOS ONE 10: e0137578.

Bai Y, Müller DB, Srinivas G, et al. (2015) Functional overlap of the *Arabidopsis* leaf and root microbiota. Nature 528: 364–369.

Bakker P, Berendsen R, Doornbos R, Wintermans P & Pieterse C (2013) The rhizosphere revisited: root microbiomics. Frontiers in Plant Science 4: 165.

Bakker PAHM, Berendsen RL, Van Pelt JA, et al. (2020) The soil-borne identity and microbiome-assisted agriculture: Looking back to the future. Molecular Plant 13: 1394–1401.

Banerjee S, Schlaeppi K & van der Heijden MGA (2018) Keystone taxa as drivers of microbiome structure and functioning. Nature Reviews Microbiology 16: 567–576.

Barret M, Morrissey JP & O’Gara F (2011) Functional genomics analysis of plant growth-promoting rhizobacterial traits involved in rhizosphere competence. Biology and Fertility of Soils 47: 729–743.

Bayer-Santos E, de Moraes Ceseti L, Farah CS & Alvarez-Martinez CE (2019) Distribution, function and regulation of Type 6 secretion systems of Xanthomonadales. Frontiers in Microbiology 10: 458221.

Belikov SI, Petrushin IS & Chernogor LI (2021) Genome Analysis of the *Janthinobacterium* sp. Strain SLB01 from the Diseased Sponge of the *Lubomirskia baicalensis*. Current Issues in Molecular Biology 43: 2220–2237.

Berendsen RL, van Verk MC, Stringlis IA, Zamioudis C, Tommassen J, Pieterse CMJ & Bakker PAHM (2015) Unearthing the genomes of plant-beneficial *Pseudomonas* model strains WCS358, WCS374 and WCS417. BMC Genomics 16: 539.

Bernal P, Llamas MA & Filloux A (2018) Type VI secretion systems in plant-associated bacteria. Environmental Microbiology 20: 1–15.

Bernal P, Allsopp LP, Filloux A & Llamas MA (2017) The *Pseudomonas putida* T6SS is a plant warden against phytopathogens. The ISME Journal 11: 972–987.

Berrios L (2021) Plant-growth-promoting *Caulobacter* strains isolated from distinct plant hosts share conserved genetic factors involved in beneficial plant–bacteria interactions. Archives of Microbiology 204: 43.

Boak EN, Kirolos S, Pan H, Pierson LS & Pierson EA (2022) The Type VI secretion systems in plant-beneficial bacteria modulate prokaryotic and eukaryotic interactions in the rhizosphere. Frontiers in Microbiology 13: 843092.

Bodelón G, Palomino C & Fernández LÁ (2013) Immunoglobulin domains in *Escherichia coli* and other enterobacteria: from pathogenesis to applications in antibody technologies. FEMS Microbiology Reviews 37: 204–250.

Borrero de Acuña JM & Bernal P (2021) Plant holobiont interactions mediated by the type VI secretion system and the membrane vesicles: promising tools for a greener agriculture. Environmental Microbiology 23: 1830–1836.

Buchfink B, Xie C & Huson DH (2015) Fast and sensitive protein alignment using DIAMOND. Nature Methods 12: 59–60.

Bulgarelli D, Garrido-Oter R, Münch Philipp C, Weiman A, Dröge J, Pan Y, McHardy Alice C & Schulze-Lefert P (2015) Structure and function of the bacterial root microbiota in wild and domesticated barley. Cell Host & Microbe 17: 392–403.

Capella-Gutiérrez S, Silla-Martínez JM & Gabaldón T (2009) trimAl: a tool for automated alignment trimming in large-scale phylogenetic analyses. Bioinformatics 25: 1972–1973.

Carrión VJ, Perez-Jaramillo J, Cordovez V, et al. (2019) Pathogen-induced activation of disease-suppressive functions in the endophytic root microbiome. Science 366: 606–612.

Castiblanco LF & Sundin GW (2016) New insights on molecular regulation of biofilm formation in plant-associated bacteria. Journal of Integrative Plant Biology 58: 362–372.

Chen I-MA, Chu K, Palaniappan K, et al. (2022) The IMG/M data management and analysis system v.7: content updates and new features. Nucleic Acids Research 51: D723–D732.

Cianciotto NP & White RC (2017) Expanding Role of Type II Secretion in Bacterial Pathogenesis and Beyond. Infection and Immunity 85: e00014–00017.

Denise R, Abby SS & Rocha EPC (2020) The Evolution of Protein Secretion Systems by Co-option and Tinkering of Cellular Machineries. Trends in Microbiology 28: 372–386.

Dias GM, de Sousa Pires A, Grilo VS, Castro MR, de Figueiredo Vilela L & Neves BC (2019) Comparative genomics of *Paraburkholderia kururiensis* and its potential in bioremediation, biofertilization, and biocontrol of plant pathogens. MicrobiologyOpen 8: e00801.

Durán D, Bernal P, Vazquez-Arias D, Blanco-Romero E, Garrido-Sanz D, Redondo-Nieto M, Rivilla R & Martín M (2021) *Pseudomonas fluorescens* F113 type VI secretion systems mediate bacterial killing and adaption to the rhizosphere microbiome. Scientific Reports 11: 5772.

Eichinger V, Nussbaumer T, Platzer A, Jehl MA, Arnold R & Rattei T (2016) EffectiveDB--updates and novel features for a better annotation of bacterial secreted proteins and Type III, IV, VI secretion systems. Nucleic Acids Res 44: D669–674.

Entila F, Han X, Mine A, Schulze-Lefert P & Tsuda K (2024) Commensal lifestyle regulated by a negative feedback loop between *Arabidopsis* ROS and the bacterial T2SS. Nature Communications 15: 456.

Estoppey A, Weisskopf L, Di Francesco E, Vallat-Michel A, Bindschedler S, Chain PS & Junier P (2022) Improved methods to assess the effect of bacteria on germination of fungal spores. FEMS Microbiology Letters 369: fnac034.

Fan E, Chauhan N, Udatha DBRKG, Leo JC & Linke D (2016) Type V secretion systems in bacteria. Microbiology Spectrum 4: 4.1.10.

Finkel OM, Salas-González I, Castrillo G, Conway JM, Law TF, Teixeira PJPL, Wilson ED, Fitzpatrick CR, Jones CD & Dangl JL (2020) A single bacterial genus maintains root growth in a complex microbiome. Nature 587: 103–108.

Franzosa EA, McIver LJ, Rahnavard G, et al. (2018) Species-level functional profiling of metagenomes and metatranscriptomes. Nature Methods 15: 962–968.

Fu L, Niu B, Zhu Z, Wu S & Li W (2012) CD-HIT: accelerated for clustering the next-generation sequencing data. Bioinformatics 28: 3150–3152.

Galán JE (2009) Common themes in the design and function of bacterial effectors. Cell Host Microbe 5: 571–579.

Gallegos-Monterrosa R & Coulthurst SJ (2021) The ecological impact of a bacterial weapon: microbial interactions and the Type VI secretion system. FEMS Microbiology Reviews 45: fuab033.

Geller AM, Pollin I, Zlotkin D, Danov A, Nachmias N, Andreopoulos WB, Shemesh K & Levy A (2021) The extracellular contractile injection system is enriched in environmental microbes and associates with numerous toxins. Nature Communications 12: 3743.

Gerlach RG & Hensel M (2007) Protein secretion systems and adhesins: The molecular armory of Gram-negative pathogens. International Journal of Medical Microbiology 297: 401–415.

Grabherr MG, Haas BJ, Yassour M, et al. (2011) Full-length transcriptome assembly from RNA-Seq data without a reference genome. Nat Biotechnol 29: 644–652.

Green ER, Mecsas J & Kudva IT (2016) Bacterial secretion systems: An overview. Microbiology Spectrum 4: 4.1.13.

Haas BJ (2023) TransDecoder V.5, https://github.com/TransDecoder/TransDecoder.

Howe AC, Jansson JK, Malfatti SA, Tringe SG, Tiedje JM & Brown CT (2014) Tackling soil diversity with the assembly of large, complex metagenomes. Proceedings of the National Academy of Sciences 111: 4904–4909.

Hu Y, Huang H, Cheng X, Shu X, White AP, Stavrinides J, Köster W, Zhu G, Zhao Z & Wang Y (2017) A global survey of bacterial type III secretion systems and their effectors. Environmental Microbiology 19: 3879–3895.

Hui X, Chen Z, Zhang J, Lu M, Cai X, Deng Y, Hu Y & Wang Y (2021) Computational prediction of secreted proteins in gram-negative bacteria. Computational and Structural Biotechnology Journal 19: 1806–1828.

Hyatt D, Chen G-L, LoCascio PF, Land ML, Larimer FW & Hauser LJ (2010) Prodigal: prokaryotic gene recognition and translation initiation site identification. BMC Bioinformatics 11: 119.

James RH, Deme JC, Hunter A, Berks BC & Lea SM (2022) Structures of the Type IX Secretion/Gliding Motility Motor from across the Phylum *Bacteroidetes*. mBio 13: e00267–00222.

Jiang F, Wang X, Wang B, Chen L, Zhao Z, Waterfield NR, Yang G & Jin Q (2016) The *Pseudomonas aeruginosa* Type VI Secretion PGAP1-like Effector Induces Host Autophagy by Activating Endoplasmic Reticulum Stress. Cell Reports 16: 1502–1509.

Johnston-Monje D, Gutiérrez JP & Lopez-Lavalle LAB (2021) Seed-transmitted bacteria and fungi dominate juvenile plant microbiomes. Frontiers in Microbiology 12: 737616.

Jones P, Binns D, Chang H-Y, et al. (2014) InterProScan 5: genome-scale protein function classification. Bioinformatics 30: 1236–1240.

Jurėnas D & Journet L (2021) Activity, delivery, and diversity of Type VI secretion effectors. Molecular Microbiology 115: 383–394.

Katoh K & Standley DM (2013) MAFFT multiple sequence alignment software version 7: improvements in performance and usability. Mol Biol Evol 30: 772–780.

Kieser S, Brown J, Zdobnov EM, Trajkovski M & McCue LA (2020) ATLAS: a Snakemake workflow for assembly, annotation, and genomic binning of metagenome sequence data. BMC Bioinformatics 21: 257.

Klockgether J & Tümmler B (2017) Recent advances in understanding *Pseudomonas aeruginosa* as a pathogen. F1000Research 6: 1261.

Kopylova E, Noé L & Touzet H (2012) SortMeRNA: fast and accurate filtering of ribosomal RNAs in metatranscriptomic data. Bioinformatics 28: 3211–3217.

Langmead B & Salzberg SL (2012) Fast gapped-read alignment with Bowtie 2. Nature Methods 9: 357–359.

Lemanceau P, Expert D, Gaymard F, Bakker PAHM & Briat JF (2009) Chapter 12 Role of Iron in Plant–Microbe Interactions. Advances in Botanical Research, Vol. 51 p. 491-549. Academic Press.

Letunic I & Bork P (2021) Interactive Tree Of Life (iTOL) v5: an online tool for phylogenetic tree display and annotation. Nucleic Acids Research 49: W293–W296.

Levy A, Salas Gonzalez I, Mittelviefhaus M, et al. (2018) Genomic features of bacterial adaptation to plants. Nature Genetics 50: 138–150.

Leyton DL, Rossiter AE & Henderson IR (2012) From self sufficiency to dependence: mechanisms and factors important for autotransporter biogenesis. Nature Reviews Microbiology 10: 213–225.

Li E, Zhang H, Jiang H, Pieterse CMJ, Jousset A, Bakker PAHM & de Jonge R (2021) Experimental-evolution-driven identification of *Arabidopsis* rhizosphere competence genes in *Pseudomonas protegens*. mBio 12: e00927–00921.

Liang J, Hoffrichter A, Brachmann A & Marín M (2018) Complete genome of *Rhizobium leguminosarum* Norway, an ineffective Lotus micro-symbiont. Standards in Genomic Sciences 13: 36.

Lin H-H, Filloux A & Lai E-M (2020) Role of Recipient Susceptibility Factors During Contact-Dependent Interbacterial Competition. Frontiers in Microbiology 11: 603652.

Liu Y, Shu X, Chen L, et al. (2023) Plant commensal type VII secretion system causes iron leakage from roots to promote colonization. Nature Microbiology 8: 1434–1449.

López JL, Fourie A, Poppeliers SWM, Pappas N, Sánchez-Gil JJ, de Jonge R & Dutilh BE (2023) Growth rate is a dominant factor predicting the rhizosphere effect. The ISME Journal 17: 1396–1405.

Love MI, Huber W & Anders S (2014) Moderated estimation of fold change and dispersion for RNA-seq data with DESeq2. Genome Biology 15: 550.

Lucke M, Correa MG & Levy A (2020) The role of secretion systems, effectors, and secondary metabolites of beneficial rhizobacteria in interactions with plants and microbes. Frontiers in Plant Science 11: 589416.

Luo Y, Frey EA, Pfuetzner RA, Creagh AL, Knoechel DG, Haynes CA, Finlay BB & Strynadka NCJ (2000) Crystal structure of enteropathogenic *Escherichia coli* intimin–receptor complex. Nature 405: 1073–1077.

Madhaiyan M, Poonguzhali S, Senthilkumar M, Pragatheswari D, Lee J-S & Lee K-C (2015) *Arachidicoccus rhizosphaerae* gen. nov., sp. nov., a plant-growth-promoting bacterium in the family Chitinophagaceae isolated from rhizosphere soil. International Journal of Systematic and Evolutionary Microbiology 65: 578–586.

McMurdie PJ & Holmes S (2013) phyloseq: An R Package for Reproducible Interactive Analysis and Graphics of Microbiome Census Data. PLOS ONE 8: e61217.

Menzel P, Ng KL & Krogh A (2016) Fast and sensitive taxonomic classification for metagenomics with Kaiju. Nature Communications 7: 11257.

Mikryukov V (2019) metagMisc: miscellaneous functions for metagenomic analysis. R-package

Minh BQ, Schmidt HA, Chernomor O, Schrempf D, Woodhams MD, von Haeseler A & Lanfear R (2020) IQ-TREE 2: New Models and Efficient Methods for Phylogenetic Inference in the Genomic Era. Molecular Biology and Evolution 37: 1530–1534.

Mistry J, Chuguransky S, Williams L, et al. (2020) Pfam: The protein families database in 2021. Nucleic Acids Research 49: D412–D419.

Nurk S, Meleshko D, Korobeynikov A & Pevzner PA (2017) metaSPAdes: a new versatile metagenomic assembler. Genome Res 27: 824–834.

Ofek-Lalzar M, Sela N, Goldman-Voronov M, Green SJ, Hadar Y & Minz D (2014) Niche and host-associated functional signatures of the root surface microbiome. Nature Communications 5: 4950.

Okazaki S, Okabe S, Higashi M, Shimoda Y, Sato S, Tabata S, Hashiguchi M, Akashi R, Göttfert M & Saeki K (2010) Identification and functional analysis of Type III effector proteins in *Mesorhizobium loti*. Molecular Plant-Microbe Interactions 23: 223–234.

Oksanen J, Blanchet FG, Kindt R, Legendre P, Minchin PR, O’hara R, Simpson GL, Solymos P, Stevens MHH & Wagner H (2013) Package ‘vegan’. Community ecology package, version 2: 1–295.

Parks DH, Chuvochina M, Rinke C, Mussig AJ, Chaumeil P-A & Hugenholtz P (2021) GTDB: an ongoing census of bacterial and archaeal diversity through a phylogenetically consistent, rank normalized and complete genome-based taxonomy. Nucleic Acids Research 50: D785–D794.

Pascale A, Proietti S, Pantelides IS & Stringlis IA (2020) Modulation of the root microbiome by plant molecules: The basis for targeted disease suppression and plant growth promotion. Frontiers in Plant Science 10: 501717.

Pfeilmeier S, Werz A, Ote M, Bortfeld-Miller M, Kirner P, Keppler A, Hemmerle L, Gäbelein CG, Pestalozzi CM & Vorholt JA (2023) Dysbiosis of a leaf microbiome is caused by enzyme secretion of opportunistic *Xanthomonas* strains. bioRxiv 2023.2005.2009.539948.

Piromyou P, Songwattana P, Boonchuen P, Nguyen HP, Manassila M, Tantanuch W, Maikhunthod B, Teamtisong K, Tittabut P & Boonkerd N (2021) The new putative Type III effector SkP48 in *Bradyrhizobium* sp. DOA9 is involved in legume nodulation. Research Square PREPRINT.

Poppeliers SWM, Sánchez-Gil JJ & de Jonge R (2023) Microbes to support plant health: understanding bioinoculant success in complex conditions. Current Opinion in Microbiology 73: 102286.

Putri GH, Anders S, Pyl PT, Pimanda JE & Zanini F (2022) Analysing high-throughput sequencing data in Python with HTSeq 2.0. Bioinformatics 38: 2943–2945.

R Core Team, 2020. R: A language and environment for statistical computing. R Foundation for Statistical Computing, Vienna, Austria. URL https://www.R-project.org/

RStudio Team, 2019. RStudio: Integrated Development for R. RStudio, Inc., Boston, MA URL http://www.rstudio.com/

Rojas-Lopez M, Zorgani MA, Kelley LA, Bailly X, Kajava AV, Henderson IR, Polticelli F, Pizza M, Rosini R & Desvaux M (2018) Identification of the Autochaperone Domain in the Type Va Secretion System (T5aSS): Prevalent Feature of Autotransporters with a β-Helical Passenger. Frontiers in Microbiology 8: 2607.

Rosier A, Medeiros FHV & Bais HP (2018) Defining plant growth promoting rhizobacteria molecular and biochemical networks in beneficial plant-microbe interactions. Plant and Soil 428: 35–55.

Russell Alistair B, Wexler Aaron G, Harding Brittany N, et al. (2014) A Type VI Secretion-Related Pathway in Bacteroidetes Mediates Interbacterial Antagonism. Cell Host & Microbe 16: 227–236.

Salomon D, Kinch LN, Trudgian DC, Guo X, Klimko JA, Grishin NV, Mirzaei H & Orth K (2014) Marker for type VI secretion system effectors. Proceedings of the National Academy of Sciences 111: 9271–9276.

Sánchez-Gil JJ, Poppeliers SWM, Vacheron J, Zhang H, Odijk B, Keel C & de Jonge R (2023) The conserved *iol* gene cluster in *Pseudomonas* is involved in rhizosphere competence. Current Biology 33: 3097–3110.e3096.

Schneijderberg M, Cheng X, Franken C, et al. (2020) Quantitative comparison between the rhizosphere effect of *Arabidopsis thaliana* and co-occurring plant species with a longer life history. The ISME Journal 14: 2433–2448.

Selten G, Stassen MJJ, De Rooij P, Berendsen RL, Stringlis I & de Jonge R (2024) Draft genome sequences of *Arabidopsis thaliana*-associated micro-organisms from Reijerscamp soil, the Netherlands [Data set]. Zenodo 10.5281/zenodo.10992416.

Seneviratne G, Weerasekara MLMAW, Seneviratne KACN, Zavahir JS, Kecskés ML & Kennedy IR (2011) Importance of Biofilm Formation in Plant Growth Promoting Rhizobacterial Action. Plant Growth and Health Promoting Bacteria,(Maheshwari DK, ed.) p. 81-95. Springer Berlin Heidelberg, Berlin, Heidelberg.

Song W, Zhuang X, Tan Y, Qi Q & Lu X (2022) The type IX secretion system: Insights into its function and connection to glycosylation in *Cytophaga hutchinsonii*. Engineering Microbiology 2: 100038.

Staehelin C & Krishnan Hari B (2015) Nodulation outer proteins: double-edged swords of symbiotic rhizobia. Biochemical Journal 470: 263–274.

Stringlis IA, Zamioudis C, Berendsen RL, Bakker PAHM & Pieterse CMJ (2019) Type III Secretion System of beneficial rhizobacteria *Pseudomonas simiae* WCS417 and *Pseudomonas defensor* WCS374. Frontiers in Microbiology 10: 467989.

Stringlis IA, Yu K, Feussner K, de Jonge R, Van Bentum S, Van Verk MC, Berendsen RL, Bakker PAHM, Feussner I & Pieterse CMJ (2018) MYB72-dependent coumarin exudation shapes root microbiome assembly to promote plant health. Proceedings of the National Academy of Sciences 115: E5213–E5222.

Suarez G, Sierra JC, Erova TE, Sha J, Horneman AJ & Chopra AK (2010) A Type VI Secretion System effector protein, VgrG1, from *Aeromonas hydrophila* that induces host cell toxicity by ADP ribosylation of actin. Journal of Bacteriology 192: 155–168.

Thomas S, Holland IB & Schmitt L (2014) The Type 1 secretion pathway — The hemolysin system and beyond. Biochimica et Biophysica Acta (BBA) - Molecular Cell Research 1843: 1629–1641.

Verster AJ, Ross BD, Radey MC, Bao Y, Goodman AL, Mougous JD & Borenstein E (2017) The landscape of Type VI Secretion across human gut microbiomes reveals its role in community composition. Cell Host & Microbe 22: 411–419.e414.

Vogel CM, Potthoff DB, Schäfer M, Barandun N & Vorholt JA (2021) Protective role of the *Arabidopsis* leaf microbiota against a bacterial pathogen. Nature Microbiology 6: 1537–1548.

von Meijenfeldt FAB, Arkhipova K, Cambuy DD, Coutinho FH & Dutilh BE (2019) Robust taxonomic classification of uncharted microbial sequences and bins with CAT and BAT. Genome Biology 20: 217.

Wallner A, Moulin L, Busset N, Rimbault I & Béna G (2021) Genetic diversity of Type 3 Secretion System in *Burkholderia s.l.* and links with plant host adaptation. Frontiers in Microbiology 12: 761215.

Wang J, Ma C, Ma S, et al. (2022) GmARP is related to the Type III effector NopAA to promote nodulation in soybean (*Glycine max*). Frontiers in Genetics 13: 889795.

Wattam AR, Davis JJ, Assaf R, et al. (2017) Improvements to PATRIC, the all-bacterial Bioinformatics Database and Analysis Resource Center. Nucleic Acids Res 45: D535–d542.

Wen T, Yuan J, He X, Lin Y, Huang Q & Shen Q (2020) Enrichment of beneficial cucumber rhizosphere microbes mediated by organic acid secretion. Horticulture Research 7: 154.

Wilhelm S, Rosenau F, Kolmar H & Jaeger K-E (2011) Autotransporters with GDSL passenger domains: Molecular physiology and biotechnological applications. ChemBioChem 12: 1476–1485.

Willett JLE, Ruhe ZC, Goulding CW, Low DA & Hayes CS (2015) Contact-dependent growth inhibition (CDI) and CdiB/CdiA two-partner secretion proteins. Journal of Molecular Biology 427: 3754–3765.

Wood DE, Lu J & Langmead B (2019) Improved metagenomic analysis with Kraken 2. Genome Biology 20: 257.

Xu J, Zhang Y, Zhang P, et al. (2018) The structure and function of the global citrus rhizosphere microbiome. Nature Communications 9: 4894.

Xu L, Dong Z, Chiniquy D, et al. (2021) Genome-resolved metagenomics reveals role of iron metabolism in drought-induced rhizosphere microbiome dynamics. Nature Communications 12: 3209.

Ye T, Zhou T, Li Q, Xu X, Fan X, Zhang L & Chen S (2020) *Cupriavidus* sp. HN-2, a Novel Quorum Quenching Bacterial Isolate, is a Potent Biocontrol Agent Against *Xanthomonas campestris* pv. *campestris*. Microorganisms 8: 45.

Yu P, He X, Baer M, et al. (2021) Plant flavones enrich rhizosphere Oxalobacteraceae to improve maize performance under nitrogen deprivation. Nature Plants 7: 481–499.

Zboralski A, Biessy A & Filion M (2022) Bridging the gap: Type III Secretion Systems in plant-beneficial bacteria. Microorganisms 10: 187.

Zeng C & Zou L (2017) An account of in silico identification tools of secreted effector proteins in bacteria and future challenges. Briefings in Bioinformatics 20: 110–129.

Zhao X, Wang Y, Shang Q, Li Y, Hao H, Zhang Y, Guo Z, Yang G, Xie Z & Wang R (2015) Collagen-like proteins (ClpA, ClpB, ClpC, and ClpD) are required for biofilm formation and adhesion to plant roots by *Bacillus amyloliquefaciens* FZB42. PLOS ONE 10: e0117414.

Zhou Y, Coventry DR, Gupta VVSR, et al. (2020) The preceding root system drives the composition and function of the rhizosphere microbiome. Genome Biology 21: 89.

